# Guidelines for evaluating the success of large carnivore reintroductions

**DOI:** 10.1101/2024.06.26.600404

**Authors:** Willem D. Briers-Louw, Peter Lindsey, Angela Gaylard, Bogdan Cristescu, Stijn Verschueren, Cole du Plessis, Marine Drouilly, Drew Bantlin, Tamar A. Kendon, Emma E.M. Evers, Caitlin J. Curry, João Almeida, David Gaynor, Alison J. Leslie, Vincent N. Naude

## Abstract

Anthropogenic impacts have led to widespread species decline and extirpation, thereby compelling a global movement to protect and regenerate biodiversity through holistic ecosystem restoration including reintroductions. Yet, despite the increasing practice of conservation-driven reintroduction efforts over the past century, peer-reviewed literature and policy providing criteria with which to evaluate reintroduction stages and efficacy remain limited. Without these comprehensive and quantifiable metrics of relative success, such drastic conservation intervention strategies cannot be objectively evaluated nor compared, hindering the advancement of restoration as a discipline. Herein, we systematically reviewed 227 large carnivore reintroductions of 14 terrestrial mammal species across 23 countries since 1930 to contextualize global efforts to date, and from these, have developed a standardized framework to evaluate reintroduction success. We further retrospectively determined the extent to which existing studies met these criteria towards identifying current knowledge gaps and guide future reintroduction efforts. Most large carnivore records were of Felidae (70%) reintroduced into ‘closed’ systems (69%) across southern Africa (70%). Our proposed framework provides a full suite of stages, indicators, and targets for reintroduction evaluation, which, when retrospectively applied to reviewed studies, indicated that at least one-third lacked sufficient information to effectively evaluate and compare reintroduction outcomes. This comprehensive and prioritized framework provides novel transparency and scalability to large carnivore reintroduction programs, which is increasingly required to secure the sustained support of impacted communities and stakeholder networks. Moreover, the incorporation of this framework into future practice and policy as an applied tool may directly benefit the recovery of at least 30 large carnivore species, while its principles may be applied more broadly across taxonomic groups for faunal rewilding and global ecosystem restoration.

## INTRODUCTION

Ecological restoration efforts have the greatest conservation value when these experiences translate directly into both adaptive management and comparable knowledge transfer (Heinen et al., 2020; Malhi et al., 2020; Seddon, 1999). Where human impacts have led to widespread fragmentation and loss of natural habitats, thousands of species are typically threatened with extinction (Ceballos et al., 2020; Tilman et al., 2017). Accumulatively, such threats have compelled a global movement to protect and regenerate biodiversity through ecological restoration (Hale et al., 2020; Jepson, 2022; Pettorelli et al., 2018). This movement is formally recognized within the United Nations Decade of Ecosystem Restoration (UNDER) initiative, which aims to proactively re-establish ecosystems to benefit both people and nature in the Anthropocene (Jepson, 2022). Despite the increasing global importance and practice of conservation-driven reintroduction efforts (i.e., restoring a species within its locally or functionally extirpated range; IUCN/SSC, 2013) over the past century, peer-reviewed literature and policy providing quantifiable criteria to comparably evaluate the relative success of such reintroductions remains limited or cursory for many threatened species (Bubac et al., 2019; Morris et al., 2021). Without traceable and quantifiable metrics of relative success (i.e., appropriate for the species, system, and conservation objective; Stepkovitch et al., 2022), such drastic conservation intervention strategies cannot be objectively evaluated, nor compared. More importantly, they cannot fully contribute to the advancement of restoration as a scientific discipline. Such transparent retrospection is essential to securing the long-term support of impacted communities and stakeholder networks, as well as guiding the sustainability of future policy, especially for large carnivores.

The substantial metabolic and ranging requirements of large carnivores drive most negative impacts of human-wildlife or conservation conflicts (Cusack et al., 2020; Redpath et al., 2015), and pose a substantial risk to many reintroduction efforts (Ripple et al., 2014). As such, metrics of relative success should be holistic (i.e., ecological, sociological, and management-related). A recent global review of 536 conservation translocations representing 54 carnivore species showed that evaluation criteria included factors relating to individual movement ecology, habitat or diet selection, and sociality, as well as population resilience, growth, and viability, supported by a suite of management practices. However, most studies lacked a standardized set of defined and evidenced reporting criteria with which to classify reintroductions (Stepkovitch et al., 2022). Consequently, existing metrics of full and partial success or failure varied by conservation objective, species, landscape, and post-release monitoring effort (Bubac et al., 2019; Morris et al., 2021). While these outcomes tend to serve site-and intervention-specific goals, they often lack the transparency and scalability of comparable benchmarks that are increasingly required by impacted communities, conservation practitioners, investors, and policymakers (Berger-Tal et al., 2020). Closing this knowledge gap in the adaptive management-science feedback mechanism thus clearly requires a complementary but practical tool for evaluating the success of large carnivore reintroductions.

Here, we address the inadequacies of current evaluation metrics by providing a comprehensive and standardized suite of definitions, stages, indicators, and quantifiable targets with which to evaluate and compare the success of large carnivore reintroduction efforts at any point throughout the intervention process. We derived these criteria from globally recorded efforts, with the goal of objectively classifying the relative success of terrestrial large carnivore reintroductions to date and provide a comparative framework towards furthering reintroduction science. We argue that sufficient evidence now exists to create such standardized metrics and that their plurality will compliment current efforts, while enhancing the overall efficacy of future intervention programs. With reintroductions becoming an increasingly important tool in the conservation of threatened species and landscape restoration (Ripple et al., 2014; Stepkovitch et al., 2022), the adaptive management framework proposed herein is directly applicable to global practice and policy governing the future preservation and recovery of at least 30 species of large carnivore.

## METHODS

We followed a three-phase approach for this study (Figure 1). Phase I comprised of an updated systematic review considering a century of large carnivore reintroductions; Phase II described a standardized framework for evaluating large carnivore reintroductions with specifically-defined stages, indicators, and quantifiable targets as derived from the updated systematic review and also presented a triage of stage-specific priority indicators and scoring for targeted evaluation given limited resources; whereafter Phase III considered a scoring system and retrospective gap analysis of reviewed reintroduction studies.

**FIGURE 1.**
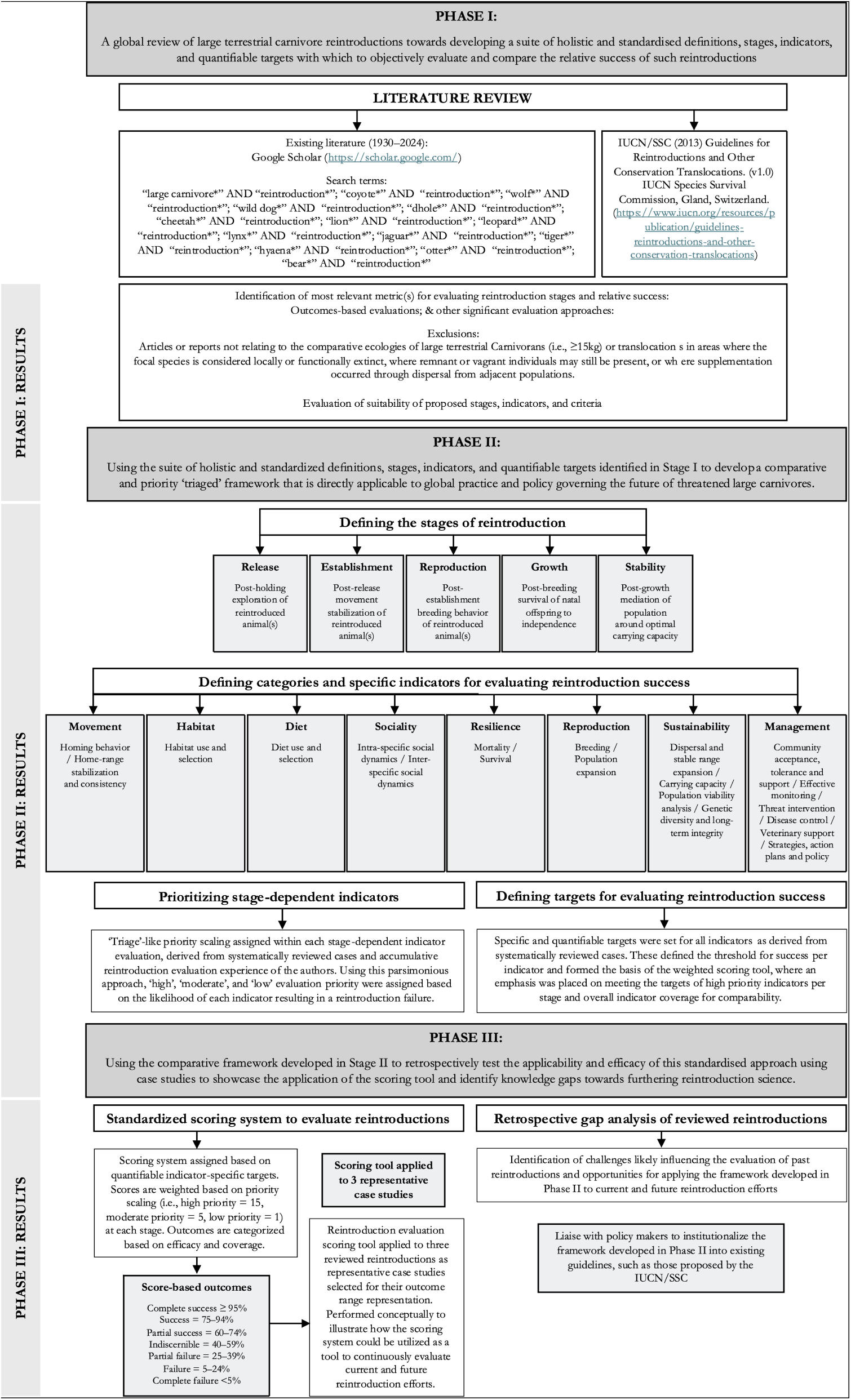
A schematic of the three-phase methodological approach. Phase I comprised of an updated systematic global review of large carnivore reintroductions. Phase II described a standardized framework for reintroductions with defined stages, indicators, and quantifiable targets as derived from the updated systematic review. Presented also, are a triage and scoring of stage-specific priority indicators for targeted evaluation given limited resources. Phase III then considered a scoring system and retrospective gap analysis of reviewed reintroduction studies.

### Phase I: Updated review

A comprehensive global review of 536 conservation-driven translocations across 54 carnivore species between the 1930s and 2022 was recently performed by Stepkovitch et al. (2022). For the purposes of this study, we updated these records, considering only the comparative ecologies of large terrestrial Carnivorans (i.e., ≥15kg; Table 1; Ripple et al., 2014), specifically identifying reintroductions (i.e., intentional translocation into areas where the focal species is considered locally or functionally extinct, and where remnant or vagrant individuals may still be present; IUCN/SSC, 2013; Stepkovitch et al., 2022). The justification for focusing on reintroductions was because these interventions inherently lack confounding interactions with resident conspecifics when animals are translocated into relatively high-density extant populations (Athreya et al., 2011; Hayward et al., 2007a; Weise et al., 2015), and arguably carry greater socio-political, economic, and ecological risks compared to augmentations (Becker et al., 2022). We conducted this systematic search through Google Scholar in April 2024, and for consistency, also used the keywords “carnivor*” or “predat*” in conjunction with “reintro*” in all searches since 1930 (Figure 1). Google Scholar was selected as this platform retrieves a greater number of records relative to alternative academic search platforms (Harzing & Alakangas, 2016). We inspected all references listed in retrieved studies, adding these to the review if relevant and not yet identified. A broader search of the grey literature (i.e., informal reports or detailed popular articles available online) was also conducted using the keywords “carnivore”, “predator”, and “reintroduction” to ensure reasonable coverage of reintroductions that have not yet been publicized.

**Table 1.**
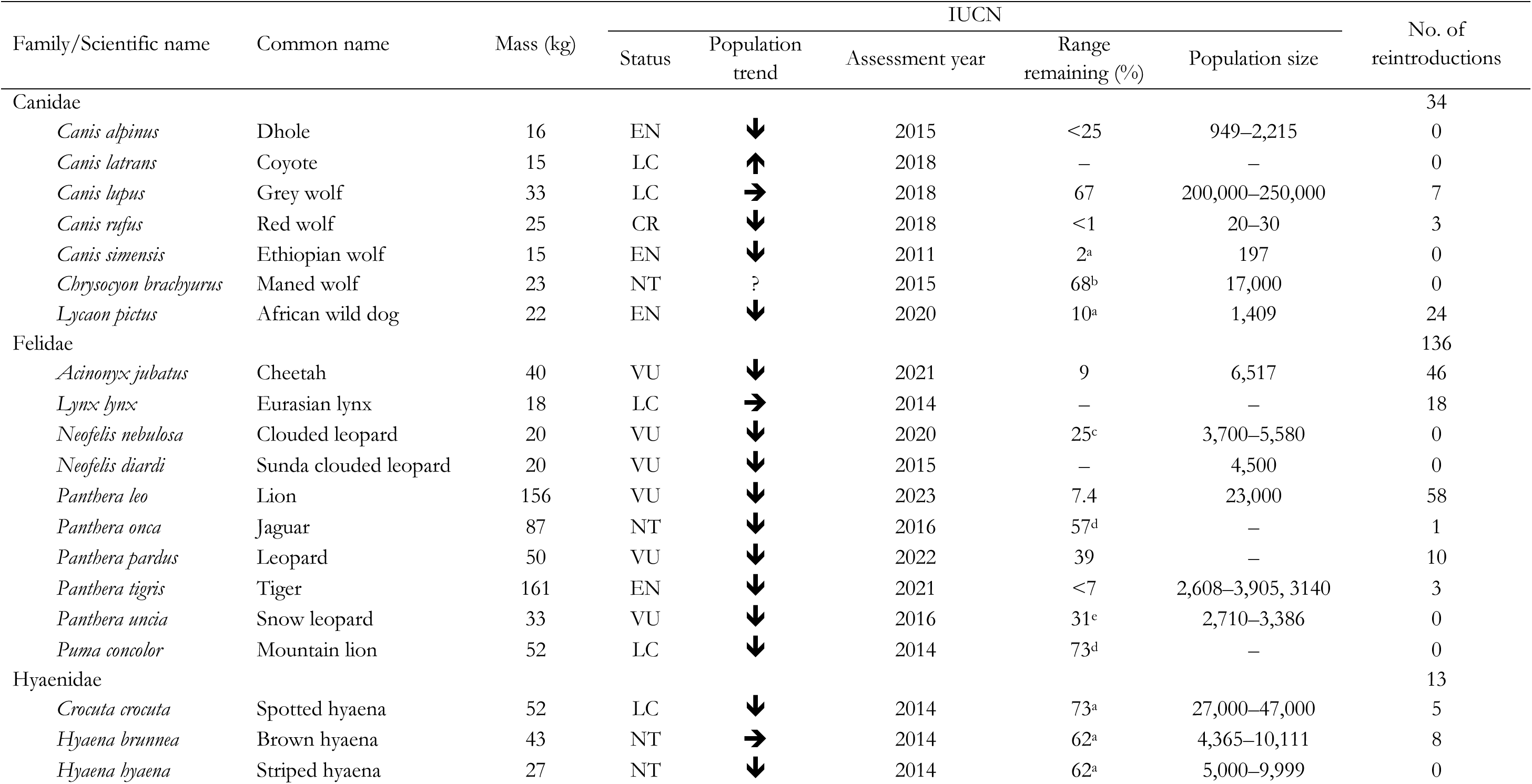
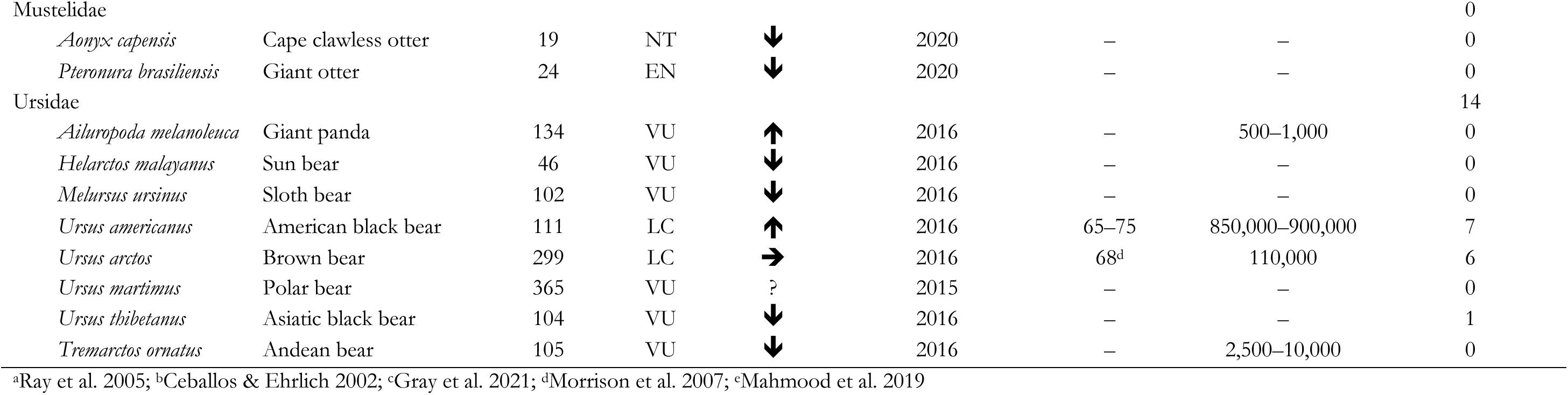
Large carnivore species, body mass (kg), species status, population trend, percent of historical range remaining, estimated number of mature individuals from wild populations, and number of reintroductions based on our review. Body masses are derived from Ripple et al. (2014) for consistency. IUCN Red List (IUCN, 2023) used for species status (LC=least concern, NT=near threatened, VU=vulnerable, EN=endangered, CR=critically endangered), trend (-f-=increasing,

To contextualize the relative success of reviewed studies, we followed the protocols of Stepkovitch et al. (2022). Reintroduction case outcomes were thus defined as: ‘failure’, ‘early’, ‘partial success’, ‘success’, or ‘unknown’ when clearly stated in-text or derived based on these classifications among reviewed studies (Table S1). A multinomial logistic regression was then employed to identify any significant differences in proportional large carnivore reintroduction outcome trends over time using the *multinom* function in the *nnet* package (Venables & Ripley 2013). Overall significance of trends was determined using type III ANOVAs implemented using the *car* package (Fox & Weisberg, 2011) and the relative effect of each outcome was plotted using the *effects* package (Fox & Hong 2009). Tukey post-hoc analyses were then carried out using the *lsmeans* package (Lenth, 2016) to test for pairwise differences in the proportion of reintroduction outcome per decade. All statistical analyses were performed in R 4.3.0 (R Core Team, 2023).

### Phase II: Standardized framework

To address several calls for an agreeable set of well-defined reporting criteria for reintroductions (Bubac et al., 2019; Morris et al., 2021; Robert et al., 2015; Seddon et al., 2007), we identified and categorized methodological consistencies from systematically reviewed studies to develop a framework for evaluating large carnivore reintroductions. This evaluation framework included a full suite of stages, indicators, and targets, along with referenced justification for all inclusions across as many species as possible (Figure 1). Terms and definitions used within our framework were derived from the Open Standards for Practice in Conservation (Conservation Measures Partnership, 2020). We did not consider pre-reintroduction feasibility stages as part of the framework, although we do strongly recommend these be conducted, as such evaluations are site-, source-and species-specific, requiring separate species-appropriate guidelines for decisions preceding reintroduction (e.g., Becker et al., 2022; Hayward et al., 2007a). Additionally, we refrain from including pre-release stages (i.e., animals translocated to the release site holding facility but not yet released) in our framework, as we argue that such events are technically classified as temporarily (semi-)captive and can thus not be evaluated until animals in holding are released.

We defined reintroduction stages (i.e., a specific and clearly defined life-history period in the reintroduction process) according to logical and unambiguous demographic, ecological and management periods with clear transitions (Miller et al., 2014). We calculated the proportion of reviewed studies which defined these stages and where necessary, logically derived these from the available information. We defined reintroduction indicators (i.e., specific, measurable variables used to track progress towards achieving reintroduction success), where we present indicators that are measurable, precise, consistent, and sensitive) based on whether they were considered to contribute to reintroduction failure if not evaluated and for convenience, indicators were grouped into broader categories. We calculated the proportion of reviewed studies which evaluated each of these indicators and where necessary, logically derived these from the available information. We defined reintroduction targets (i.e., specific, quantifiable goals set for each indicator) for success for each indicator based on reviewed studies and where necessary, logically derived these from the available information.

To account for any financial, resource, and expertise limitations often experienced at various stages by reintroduction programs (Bubac et al., 2019; Stepkovitch et al., 2022), we assigned ‘triage’-like priority scaling within each stage-dependent indicator evaluation. The evidence for such priority determination was derived from the systematically reviewed cases and the accumulative reintroduction evaluation experience of the authors. Using this parsimonious approach, we assigned ‘high’, ‘moderate’, and ‘low’ evaluation priority based on the likelihood of this indicator resulting in a reintroduction failure if unevaluated or the target is not met at that specific stage of the reintroduction process. This stage-based indicator and target framework should be considered as a sliding scale of transitional priorities (i.e., reintroduction programs can be evaluated at any time throughout the process and can both progress and regress across stages based on these criteria).

### Phase III: Scoring and gap analyses

Building on this priority framework, we developed a pragmatic scoring system for quantifying evaluation efforts (i.e., whether there was sufficient evidence to determine an outcome) and outcomes (i.e., whether priority targets were met), as well as for comparison within and between studies (i.e., tracking monitoring effort and intervention efficacy over time or between reintroductions at similar stages). Given that current definitions of reintroduction success or failure are limited by time (Seddon, 1999), we propose two categories for continuous applicability across various reintroduction stages. The first being ‘relative success’ (i.e., stage-based indicator evaluation, for programs which have not yet successfully progressed through all reintroduction stages) and thereafter ‘overall success’ (i.e., programs which successfully progressed through all reintroduction stages). Where we argue that reintroduction programs attaining ‘overall success’ or population stability over time, are marked by a ‘hand-over’ to management, after which the reintroduction program should cease, and the population be considered functionally established. While it would be inappropriate to apply this scoring system to the reviewed reintroduction cases in this study, given that these have collectively been used to develop our suite of indicators, we did conceptually apply such scoring to three representative case studies, illustrating how such stage-depended priority scoring could be used as a tool to continuously evaluate current and future reintroduction efforts.

To identify mismatches in the evaluation of indicators relative to our stage-based priority scaling framework, we carried out a retrospective knowledge gap analysis on all reviewed reintroduction cases. We calculated the proportion of reviewed studies with sufficient evidence to evaluate these stage-specific indicators, which we consider an important step towards improved contextual understanding and comparability within and between large carnivore reintroduction programs. Ultimately, this gap analysis was not aimed at re-evaluating reintroduction outcomes or criticizing past efforts. Rather, we use it to highlight inconsistencies of past reintroduction evaluation across studies and thus assess the overall value of this comprehensive and standardized evaluation framework.

## RESULTS

### Phase I: Updated review

A total of 227 large carnivore reintroduction events were recorded in the updated systematic review (Table S2), representing at least 2,212 reported individuals as 15 species over 146 sites across 23 countries between 1930 and 2024 (Figure 2). Most of these reintroductions took place in Africa (i.e., 76%, of which 70% occurred in southern Africa), followed by Europe (12%), North America (6%), Asia (45), and South America (1%). The majority of reintroductions were into ‘closed’ or functionally isolated (e.g., impermeably fenced; 69%) rather than into ‘open’ or potentially interconnected systems (31%). Reintroductions comprised of 15 focal species (i.e., 50%) from eligible carnivore families (Felidae, 70%; Canidae, 17%; Hyaenidae, 8%; Ursidae, 6%). Of these species, 47% represented ‘Threatened’ categories (i.e., IUCN Red List of Threatened Species status: ‘Vulnerable’, ‘Endangered’, or ‘Critically Endangered’), while 60% of these species are experiencing global population declines (IUCN, 2023). Since the 1930s, reintroductions have increased by an average of 20 reintroductions per decade (Figure 2), where the average number of individuals per reintroduction was 12.72 ± 1.71 [SE]. Reported reintroduction outcomes across all cases were 34% early, 29% success, 19% failure, 14% unknown/not classified, and 4% partial success. While it would be assumed that the relatively high proportion of early outcomes were attributed to more recent reintroductions, the average time between first year of reintroduction and study publication date was 4.68 ± 0.52 years (x ∼ 4 years) for these particular cases. Given that successful reintroductions were reported as early as two years post-release, most early studies would generally have had sufficient time to evaluate reintroductions, while successful and failed reintroductions were only determined after an average of 14.86 ± 1.56 and 21.34 ± 2.28 years respectively. Large carnivore reintroduction outcome trends differed significantly by decade (χ^2^ = 103.16, df = 4, *P* < 0.001), indicating that while proportional failure has declined significantly (62% to 6%, *P* < 0.05) since 1970, there has been no significant (29% vs 24%, *P* = 0.14) improvement in the proportion of successful outcomes, with unknown and early outcome proportions increasing over time.

**FIGURE 2.**
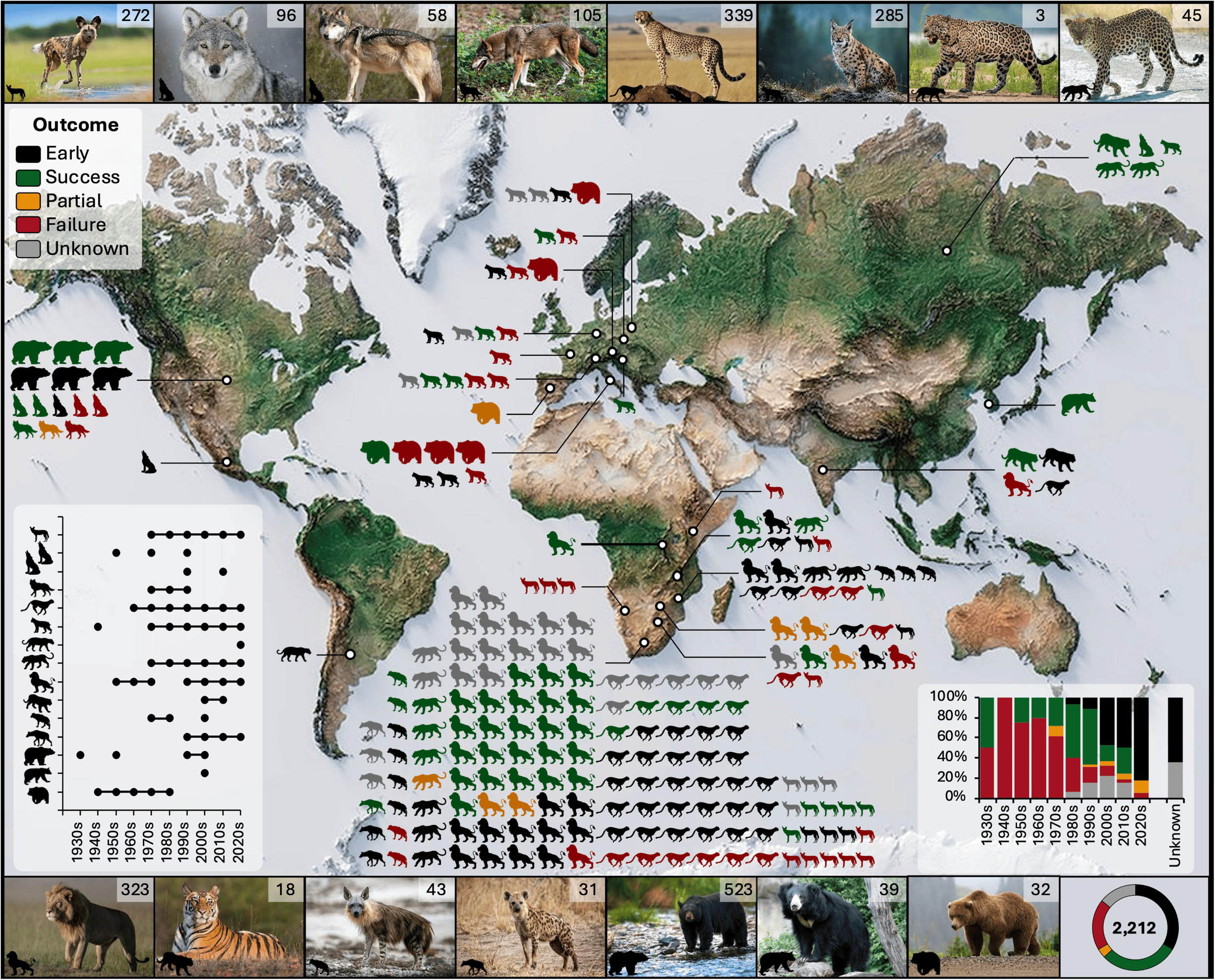
A map displaying all terrestrial, large carnivore (>15kg) reintroductions (*n* = 227) across the globe with proportional self-defined reintroductions outcomes indicated per species (i.e., black = ‘early’, green = ‘success’, yellow = ‘partial’, red = ‘failure’, and grey = ‘unknown’). Insets show reintroductions per species by decade (bottom left) and proportional outcomes for all large carnivores by decade (bottom right). The total number of individuals reintroduced per species are represented by the number overlayed on each species image (top and bottom ribbons), with the total number of individuals and their proportional outcomes indicated in the pie chart.

### Phase II: Standardized framework

#### Reintroduction stages and transitions

Evaluation of reintroduction success is a multi-stage process (Miller et al., 2014). We argue for five core stages (Table 2) in the post-release evaluation of large carnivore reintroductions: 1) release (i.e., post-holding exploration), 2) establishment (i.e., post-release movement stabilization), 3) breeding (i.e., post-establishment breeding behavior), 4) growth (i.e., post-breeding survival of natal offspring to independence), and 5) stability (i.e., post-growth mediation of population around carrying capacity). Reintroduction stages were determined from 74% of reviewed studies, while no stages could be identified for 26% of cases (i.e., unknown). Of all the reintroductions, 51% progressed from the release stage to establishment, 48% reached breeding, 31% reached growth and only 8% reached stability stage.

**TABLE 2.**
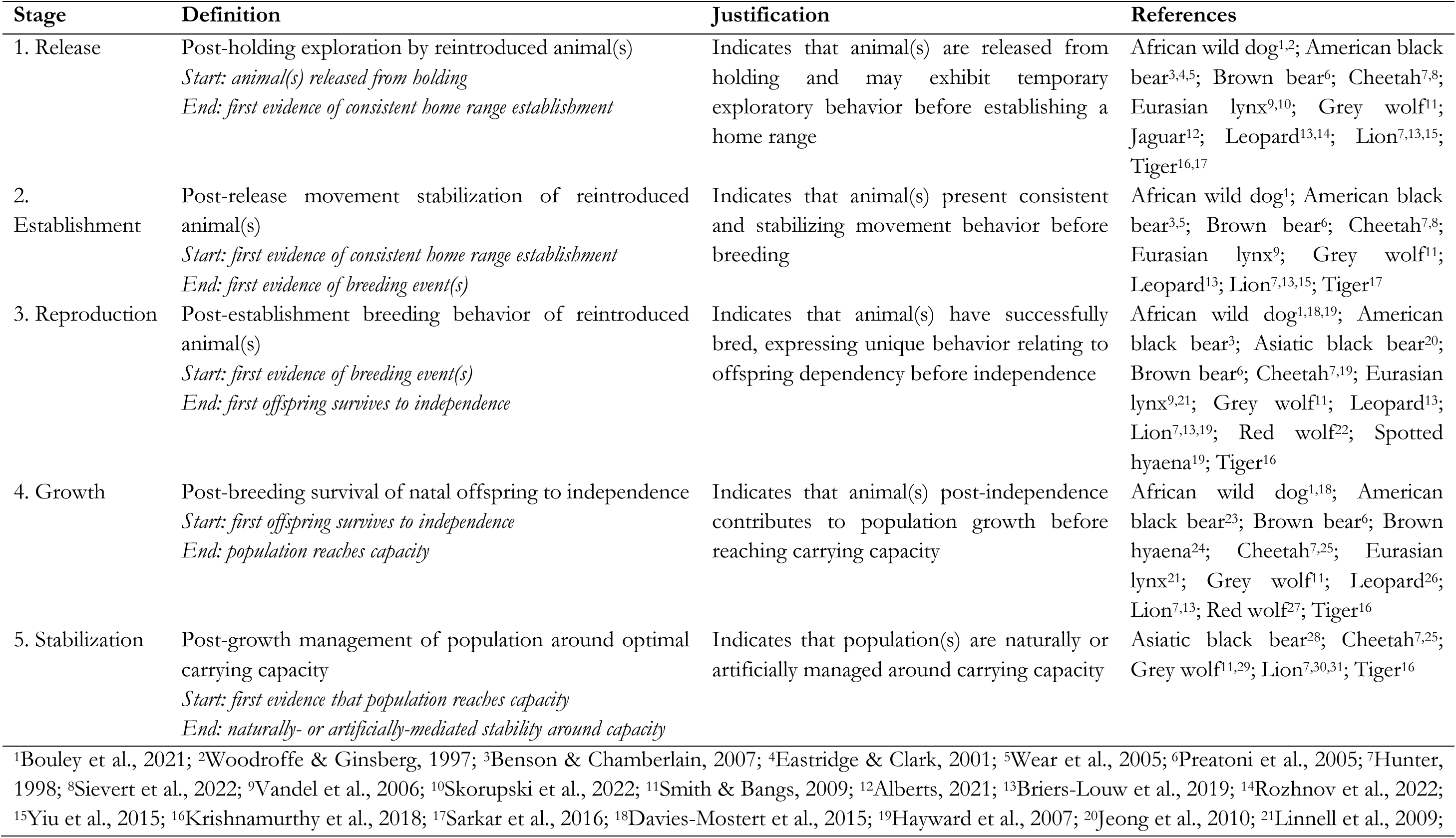
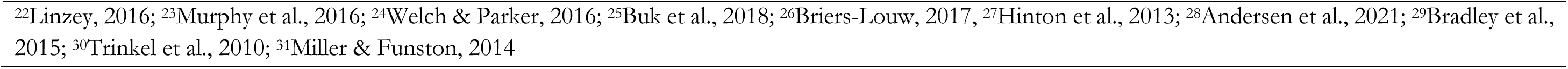
Proposed large carnivore reintroduction stages. Indicated are the stages, definitions, justifications and supporting references.

| Stage | Definition | Justification | References |
| --- | --- | --- | --- |
| 1. Release | Post-holding exploration by reintroduced animal(s)<br>*Start: animal(s) released from holding*<br>*End: first evidence of consistent home range establishment* | Indicates that animal(s) are released from holding and may exhibit temporary exploratory behavior before establishing a home range | African wild dog<sup>1,2</sup>; American black bear<sup>3,4,5</sup>; Brown bear<sup>6</sup>; Cheetah<sup>7,8</sup>; Eurasian lynx<sup>9,10</sup>; Grey wolf<sup>11</sup>; Jaguar<sup>12</sup>; Leopard<sup>13,14</sup>; Lion<sup>7,13,15</sup>; Tiger<sup>16,17</sup> |
| 2. Establishment | Post-release movement stabilization of reintroduced animal(s)<br>*Start: first evidence of consistent home range establishment*<br>*End: first evidence of breeding event(s)* | Indicates that animal(s) present consistent and stabilizing movement behavior before breeding | African wild dog<sup>1</sup>; American black bear<sup>3,5</sup>; Brown bear<sup>6</sup>; Cheetah<sup>7,8</sup>; Eurasian lynx<sup>9</sup>; Grey wolf<sup>11</sup>; Leopard<sup>13</sup>; Lion<sup>7,13,15</sup>; Tiger<sup>17</sup> |
| 3. Reproduction | Post-establishment breeding behavior of reintroduced animal(s)<br>*Start: first evidence of breeding event(s)*<br>*End: first offspring survives to independence* | Indicates that animal(s) have successfully bred, expressing unique behavior relating to offspring dependency before independence | African wild dog<sup>1,18,19</sup>; American black bear<sup>3</sup>; Asiatic black bear<sup>20</sup>; Brown bear<sup>6</sup>; Cheetah<sup>7,19</sup>; Eurasian lynx<sup>9,21</sup>; Grey wolf<sup>11</sup>; Leopard<sup>13</sup>; Lion<sup>7,13,19</sup>; Red wolf<sup>22</sup>; Spotted hyaena<sup>19</sup>; Tiger<sup>16</sup> |
| 4. Growth | Post-breeding survival of natal offspring to independence<br>*Start: first offspring survives to independence*<br>*End: population reaches capacity* | Indicates that animal(s) post-independence contributes to population growth before reaching carrying capacity | African wild dog<sup>1,18</sup>; American black bear<sup>23</sup>; Brown bear<sup>6</sup>; Brown hyaena<sup>24</sup>; Cheetah<sup>7,25</sup>; Eurasian lynx<sup>21</sup>; Grey wolf<sup>11</sup>; Leopard<sup>26</sup>; Lion<sup>7,13</sup>; Red wolf<sup>27</sup>; Tiger<sup>16</sup> |
| 5. Stabilization | Post-growth management of population around optimal carrying capacity<br>*Start: first evidence that population reaches capacity*<br>*End: naturally- or artificially-mediated stability around capacity* | Indicates that population(s) are naturally or artificially managed around carrying capacity | Asiatic black bear<sup>28</sup>; Cheetah<sup>7,25</sup>; Grey wolf<sup>11,29</sup>; Lion<sup>7,30,31</sup>; Tiger<sup>16</sup> |

<sup>1</sup>Bouley et al., 2021; <sup>2</sup>Woodroffe & Ginsberg, 1997; <sup>3</sup>Benson & Chamberlain, 2007; <sup>4</sup>Eastridge & Clark, 2001; <sup>5</sup>Wear et al., 2005; <sup>6</sup>Preatoni et al., 2005; <sup>7</sup>Hunter, 1998; <sup>8</sup>Sievert et al., 2022; <sup>9</sup>Vandel et al., 2006; <sup>10</sup>Skorupski et al., 2022; <sup>11</sup>Smith & Bangs, 2009; <sup>12</sup>Alberts, 2021; <sup>13</sup>Briers-Louw et al., 2019; <sup>14</sup>Rozhnov et al., 2022; <sup>15</sup>Yiu et al., 2015; <sup>16</sup>Krishnamurthy et al., 2018; <sup>17</sup>Sarkar et al., 2016; <sup>18</sup>Davies-Mostert et al., 2015; <sup>19</sup>Hayward et al., 2007; <sup>20</sup>Jeong et al., 2010; <sup>21</sup>Linnell et al., 2009;
<sup>22</sup>Linzey, 2016; <sup>23</sup>Murphy et al., 2016; <sup>24</sup>Welch & Parker, 2016; <sup>25</sup>Buk et al., 2018; <sup>26</sup>Briers-Louw, 2017, <sup>27</sup>Hinton et al., 2013; <sup>28</sup>Andersen et al., 2021; <sup>29</sup>Bradley et al., 2015; <sup>30</sup>Trinkel et al., 2010; <sup>31</sup>Miller & Funston, 2014

#### Reintroduction indicators

Past reintroduction evaluations generally lacked well-defined metrics and where indicators were applied, many were inconsistent and highly variable (Table S2). In the interest of clarity, replicability, and comparability across large carnivore reintroductions, we identify 20 review-derived quantifiable definitions of indicators for evaluating reintroduction success (Table 3) which align with IUCN/SSC (2013) guidelines. Overlooking any of these aspects could lead to reintroduction failure, thus we highly recommend that current and future reintroduction programs invest effort in assessing these indicators. The importance of indicator-based evaluation categories is evidenced through key case studies presented below, with associated percentages representing the proportion of indicators evaluated by all reviewed studies.

**TABLE 3.**
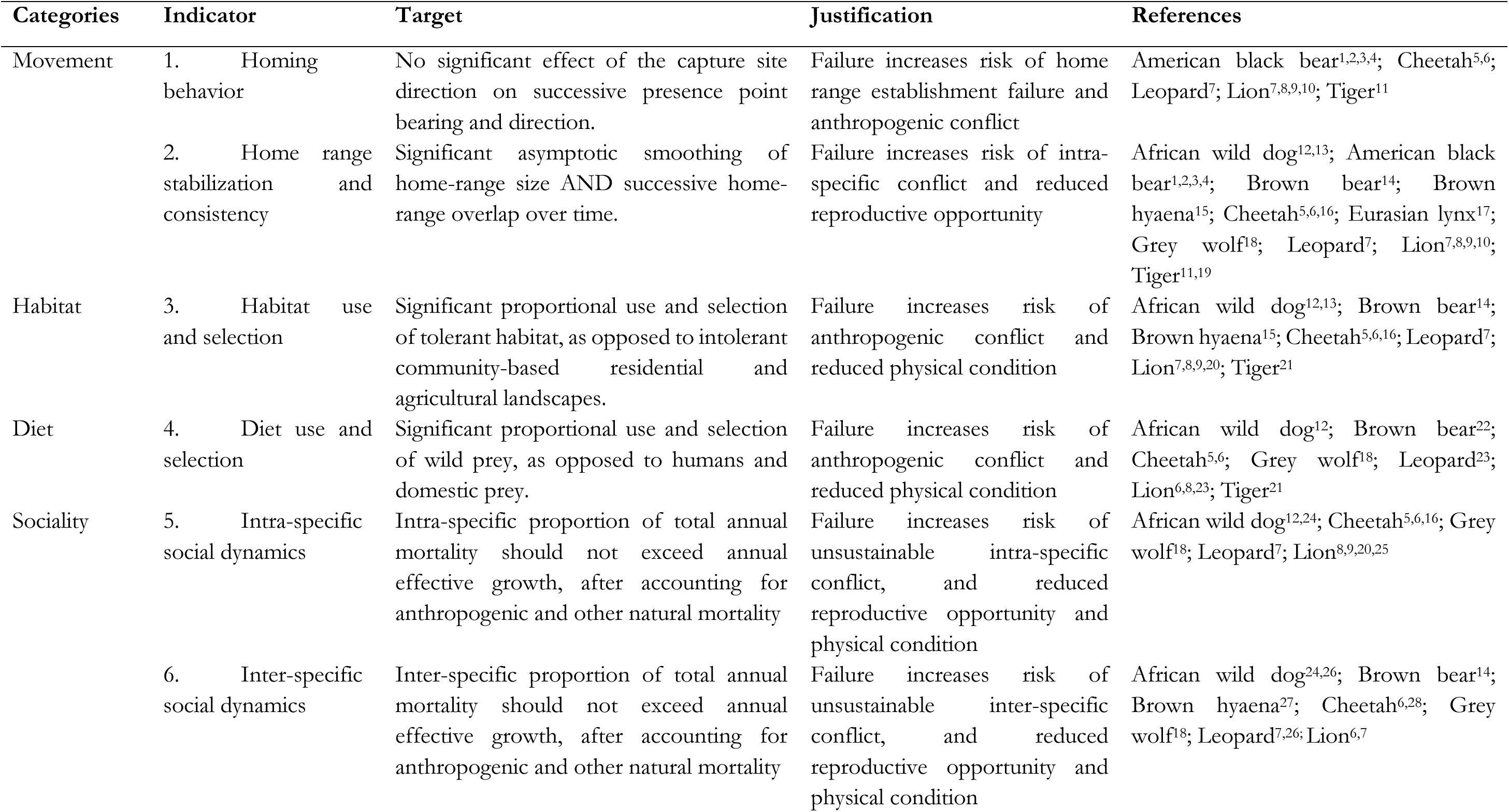
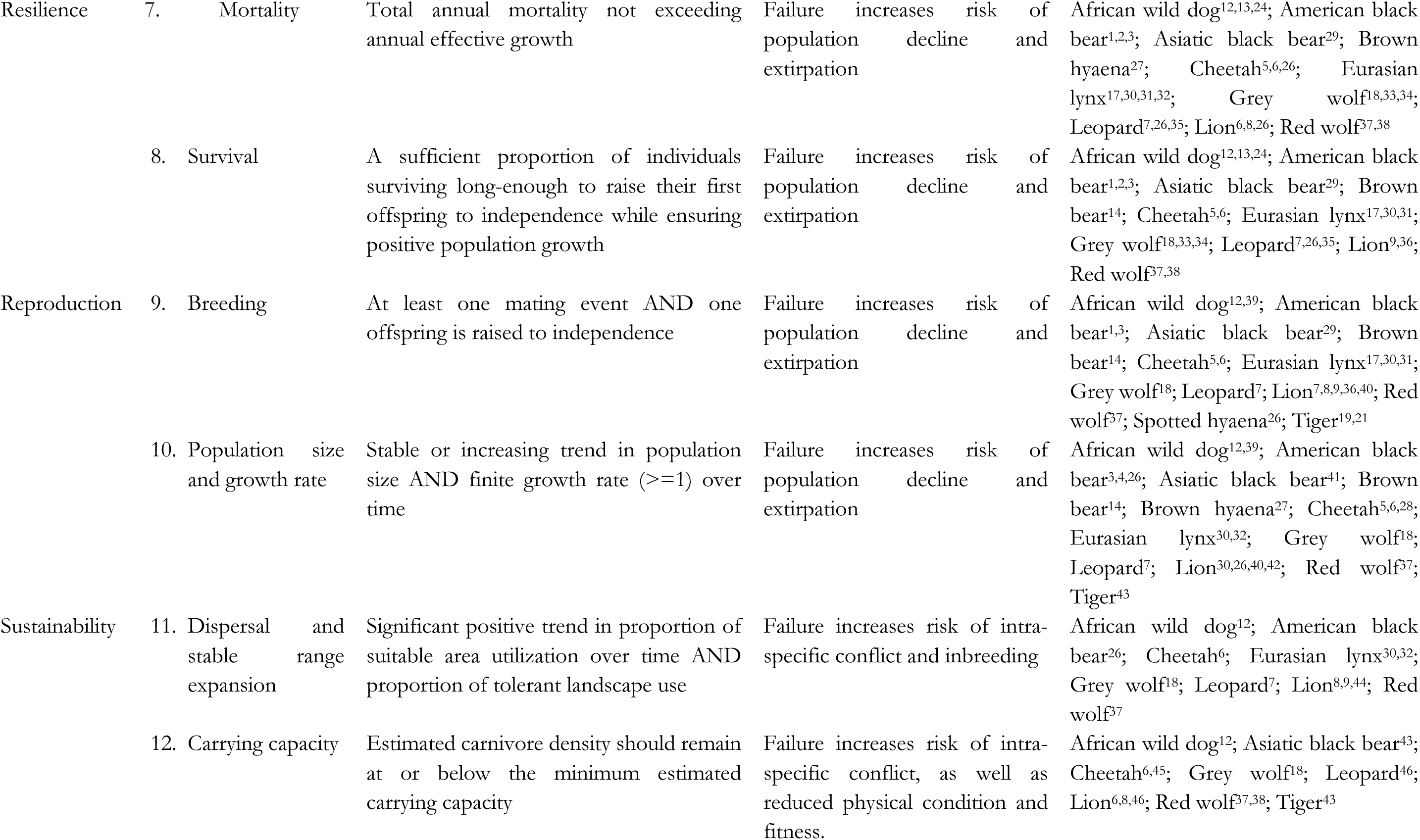
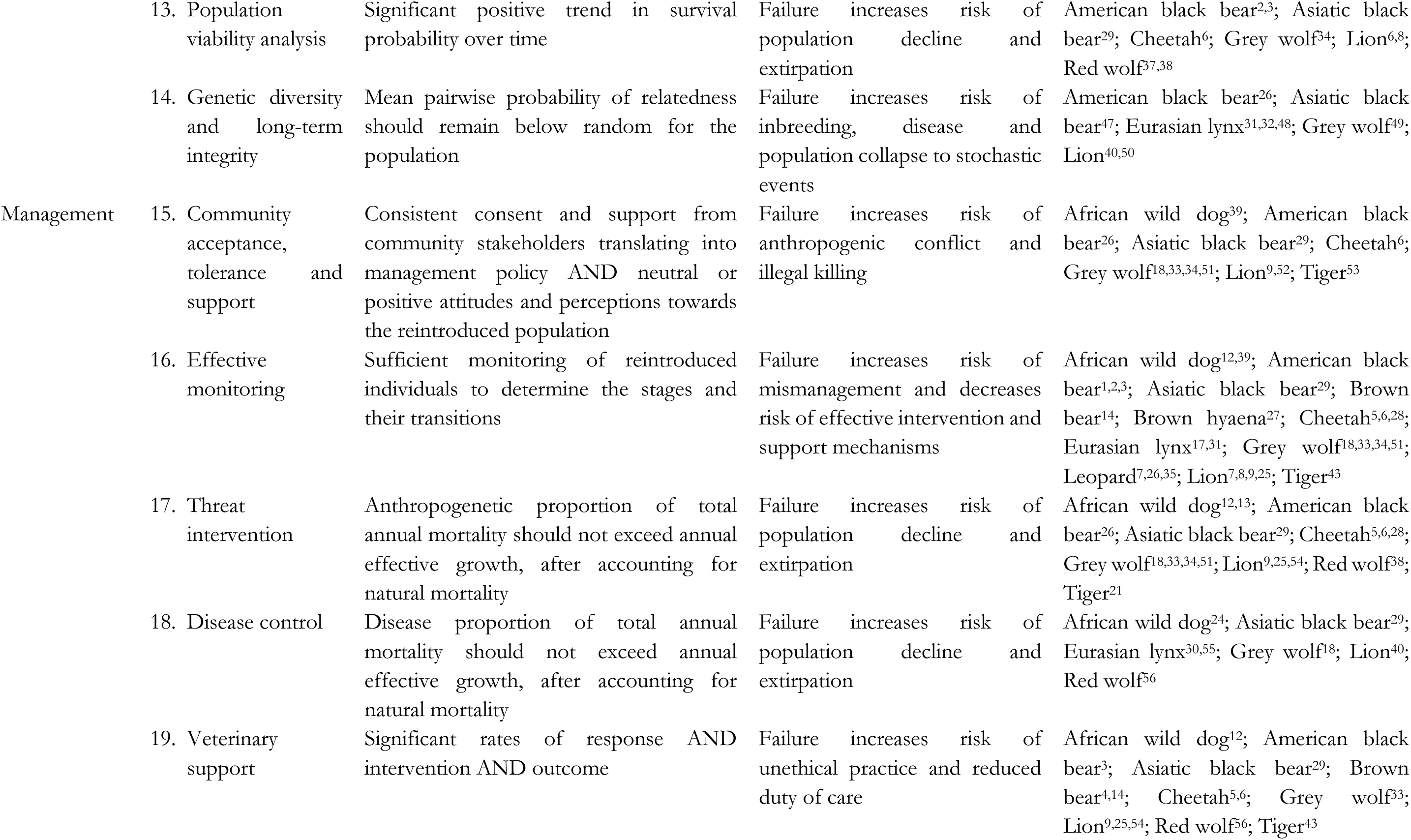
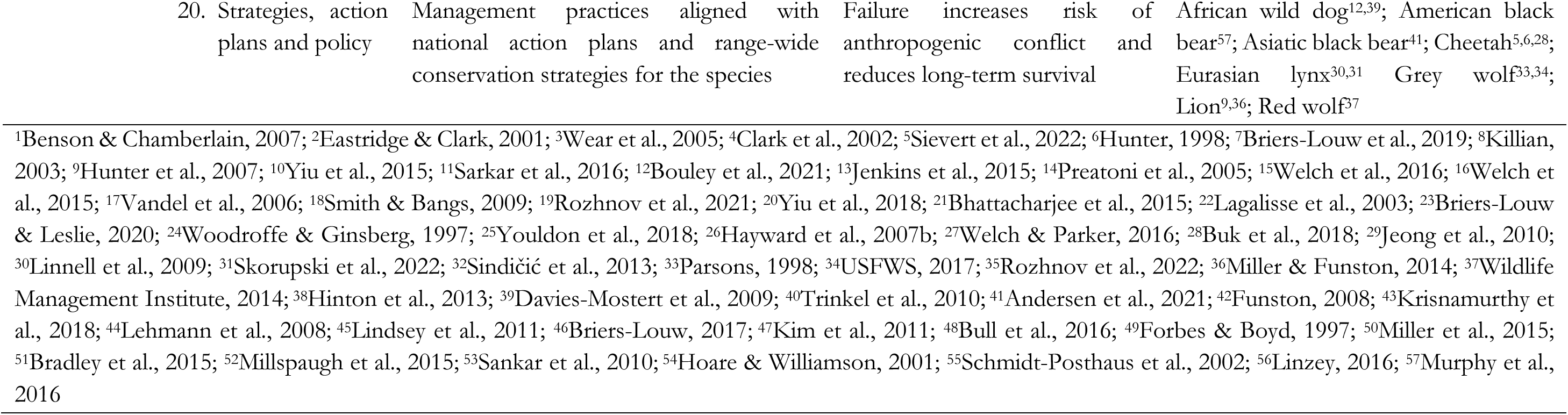
Proposed large carnivore reintroduction evaluation indicators, including categories, indicators, targets, justifications (i.e., primary consequence of failing to meet the indicated targets) and supporting references.

Movement – Reviewed studies considered two primary post-release movement indicators: homing (7%) and home range stabilization and consistency (16%), with the expectation that reintroduced large carnivores would display release site fidelity. Homing behavior proved a major obstacle for American black bear (*Ursus americanus*) reintroductions in North America (Eastridge & Clark, 2001) and was attributed to the failure of a cheetah (*Acinonyx jubatus*) reintroduction in Zambia (Phiri, 1996). However, adaptive release methods (e.g., temporary pre-release holding facilities) and large translocation distances have reduced homing and improved founder survival (Briers-Louw et al., 2019; Clark et al., 2002). While individual home range stabilization and consistency were evaluated in the reintroduction success of several large carnivores (Briers-Louw et al., 2019; Hunter et al., 2007; Preatoni et al., 2005; Rozhnov et al., 2021; Sarkar et al., 2016). The lack of range stability and consistency may also result in Allee effects as documented in grey wolves (*Canis lupus*; Hurford et al., 2006) and African wild dogs (*Lycaon pictus*; Gusset et al., 2006), or mortality during exploratory movements (Hunter et al., 2007; Preatoni et al., 2005). Given their potential contribution to reintroduction failure, we consider these indicators as important, particularly during the release and establishment stages of a reintroduction.

Habitat – Landscape use was evaluated in 11% of reviewed studies, with the expectation that large carnivores would utilize natural habitat as opposed to anthropogenically-dominated habitat. Consistent use of community land contributed to the failure of African wild dog reintroductions in South Africa, and consequent negative attitudes of landowners has likely ruled out future reintroductions of the species in the area (Davies-Mostert et al., 2009). Additionally, the lack of sufficient natural habitat and potential for human-wildlife conflict may threaten reintroduction efforts of tigers (*Panthera tigris*) in Sariska Tiger Reserve in India (Sankar et al., 2010). Given the potential impact of habitat use and preference on reintroduction failure, either through mortality or negative stakeholder attitudes, we consider this an important indicator, particularly during the release, establishment and reproduction stages of a reintroduction.

Diet – Dietary intake was evaluated in 14% of reviewed studies, typically with the expectation that large carnivores will consume wild prey as opposed to livestock. Studies found that tigers reintroduced into Sariska and Panna Tiger Reserves in India consumed high proportions of livestock which may jeopardize restoration efforts if left unaddressed (Kolipaka et al., 2017; Sankar et al., 2010). Furthermore, while the impact of livestock depredation by grey wolves on the industry appears largely negligible in the Northwestern U.S., substantial and excessive livestock killing highlighted the need for transparent engagement and efficient compensation to maintain landowner tolerance levels (Muhly & Musiana, 2009). Evaluating diet is important in reducing human-wildlife conflict, and is thus considered an important indicator, particularly during the release, establishment and reproduction stages of a reintroduction.

Sociality – Reviewed studies evaluated two sociality indicators: intra-specific (19%) and inter-specific social dynamics (16%), with the expectation that reintroduced large carnivores should display natural social stability and avoid unnaturally lethal levels of negative interaction (e.g., competition or predation). This lack of social cohesion may increase post-release roaming, dispersal, and mortality, as highlighted in the reintroductions of African wild dogs (Gusset et al., 2006) and lions (Youldon et al., 2016). Social instability may also result in slow recolonization rates and establishment of breeding pairs due to increased difficulty in finding mates at low densities, as observed in grey wolves (Hurford et al., 2006). Inter-specific social dynamics may directly or indirectly impact reintroduced large carnivores through interference or exploitative competition respectively. For example, lions alone contributed to over 30% of total cheetah mortalities across metapopulation reserves in South Africa (Buk et al., 2018) and played a significant role in the failure of African wild dog reintroductions in Etosha National Park, Namibia (Scheepers & Venzke, 1995). More subtly, reintroduced large carnivore fitness and survivability may also be influenced by other species, for example through habitat and range use (du Preez et al., 2015) and dietary ecology (Briers-Louw & Leslie, 2020; Cornhill & Kerley, 2020). Both sociality indicators are considered important, particularly during the release, establishment, and reproduction stages of a reintroduction.

Resilience – Reviewed studies evaluated mortality (37%) and survival (39%) of reintroduced large carnivores with the expectation that annual mortality should not exceed annual growth rate and that sufficient individuals survive to contribute to population growth. Snaring by-catch, an increasingly significant threat to large carnivores, caused high mortality and led to the extirpation of reintroduced African wild dogs in a South African reserve (Jenkins et al., 2015), while retaliatory killing contributed to grey wolf mortalities in North America prompting the implementation of adaptive mitigation measures (Smith & Bangs, 2009). Illegal hunting was the major driver suppressing the Eurasian lynx (*Lynx lynx*) population of central Europe, which established and increased, but later declined and stagnated; this threat was highlighted as the highest priority to address to facilitate recover (Heurich et al., 2018). Poaching was also responsible for 41% of red wolf (*Canis rufus*) pair disbandment and 34% decline in annual preservation rates of breeding pairs, threatening restoration efforts of this critically endangered Canid (Hinton et al., 2015). While a lack of founder survival resulted in the failed reintroductions of African wild dogs in southern Africa (e.g., Woodroffe & Ginsberg, 1997) and brown bears (*Ursus arctos*) in Europe (e.g., Clark et al., 2002). We consider mortality and survival important indicators, particularly during the release, establishment, and reproduction stages of a reintroduction.

Reproduction – Reviewed studies evaluated breeding (37%) and population growth (30%), with the expectation that reintroduced large carnivores should produce offspring and raise them to independence towards supporting a stable or increasing population trend. Evaluating post-release reproductive events was important to understand breeding success of grey wolves in North America (Smith & Bangs, 2009), and brown bears in Italy (Tosi et al., 2015). Additionally, reproductive rates obtained from South African metapopulation studies provide important comparative benchmarks for reintroduced African large carnivores in closed systems (e.g., Davies-Mostert et al., 2015). However, a lack of breeding or successfully raising offspring to independence may result in reintroduction failure as observed for red wolves in North America (Linzey, 2016). Population growth rates are well documented in small, closed reserves in southern Africa, particularly for lions (Miller & Funston, 2014), cheetahs (Buk et al., 2018) and African wild dogs (Davies-Mostert et al., 2015). Population assessments of reintroduced species may prove important in determining extirpated populations (e.g., cheetah; Purchase, 2007), fluctuating populations (e.g., Eurasian lynx; Heurich et al., 2018) and rapidly increasing populations (e.g., brown hyaenas *Hyaena brunnea*; Welch & Parker, 2016). We consider breeding an important indicator during the reproduction stage, and population growth an important indicator during the growth and stability stages.

Sustainability – Reviewed studies evaluated four indicators for viability: dispersal and stable range expansion (15%), carrying capacity (21%), population viability (5%) and genetic diversity (15%). Despite population increases, range expansion may be limited due to environmental or anthropogenic factors, as observed in Poland where reintroduced Eurasian lynx failed to display range expansion due to anthropogenic pressures (Skorupski et al., 2022). The lack of natural dispersal and range expansion, due to a fence, resulted in inbreeding, poor body condition and lower survival in a reintroduced lion population in Hluhluwe-iMfolozi Park, South Africa, which was later addressed through subsequent supplementation (Trinkel et al., 2008). In contrast, dispersal and range expansion may be successful, for example brown bears expanded into various countries in central Europe (Tosi et al., 2015) and grey wolf expanded across the North American Midwest (Jimenez et al., 2017). Predicting large carnivore carrying capacities based on prey and habitat availability provide fundamental recovery targets and a basis for adaptive species management. Of particular concern are populations exceeding carrying capacity which may cause ecological imbalance due to impacts on prey or competitor populations (Buk et al., 2018; Davies-Mostert et al., 2009; Lindsey et al., 2011), and exert negative human-wildlife impacts. Genetic analyses have also helped to determine the viability of reintroduced American black bear (Murphy et al., 2016) and grey wolf (von Holdt et al., 2008) populations. However, a study found genome-wide diversity loss in several reintroduced Eurasian lynx populations across central Europe (Bull et al., 2016), which demonstrates that reintroduced populations may require active genetic management when there is a lack of connectivity with other viable populations. Similarly, genetic analyses revealed that augmentation was required for genetic rescue in lions (Trinkel et al., 2008) and red wolves (Hedrick & Fredrickson, 2008). Population viability analyses can simulate potential future scenarios of reintroduced populations and provide a quantitative foundation for assessing the effect of alternative management strategies (Seddon et al., 2007). Heurich et al. (2018) used probabilistic extinction modelling to evaluate Eurasian lynx status in central Europe and determined that local extinction was likely if anthropogenic mortality was not addressed. Population viability was similarly assessed for reintroduced American black bears in North America, indicating a very low extinction risk owing to founder population size and demographic parameters (Wear et al., 2005). These indicators are considered important, particularly during the growth and stability stages of a reintroduction.

Management – Reviewed studies evaluated five indicators for management: community acceptance, tolerance and support (16%), effective monitoring (60%), threat intervention (18%), disease control (11%), veterinary support (22%), and strategies, action plans and policy (46%). Evaluating attitudes and perceptions of impacted communities before and after reintroductions may improve our understanding of tolerance levels and guide species management. For example, the clandestine reintroduction of Eurasian lynx in Switzerland, while supported by the urban populus, was opposed by rural communities and subsequent illegal hunting resulted in dramatic population reductions (Heurich et al., 2018). Attitudes towards expanding tiger populations in India varied; local communities mostly held negative attitudes towards tigers in Panna Tiger Reserve, unless compensated for livestock losses; while those in Sariska Tiger Reserve held mostly positive attitudes due to their belief system and a more coexistence-orientated lifestyle (Malviya et al., 2022). A pre-requisite for any reintroduction is to control threats which caused the initial extinction (IUCN/SSC, 2013); however, ongoing post-release intervention is also required. For example, inadequate protection efforts to limit sustained poaching resulted in the decline of Eurasian lynx in Switzerland (Arlerttaz et al., 2021) and extirpation of African wild dogs in a South African reserve (Jenkins et al., 2015). Disease outbreak control was highlighted as an important consideration in large carnivore reintroductions. For example, grey wolves reintroduced into the Yellowstone National Park in North America experienced a population crash following outbreaks of canine distemper (Smith & Bangs, 2009). Disease also contributed to 18% of reintroduced Eurasian lynx mortalities in Switzerland (Schmidt-Posthaus et al., 2002), and the extirpation of reintroduced African wild dogs in Etosha National Park in Namibia (Scheepers & Venzke, 1995) and Tswalu Kalahari Reserve in South Africa (Davies-Mostert et al., 2009). Access to veterinary support can aid rapid intervention and minimize mortality, as observed with the rescue of key Asiatic black bear (*Ursus thibetanus*) individuals (Jeong et al., 2021), while retrieval of escaped cheetah from South African reserves prevented both individual mortality and community disengagement (Buk et al., 2018). Although post-release monitoring of reintroduced carnivores is key and can assist other restoration efforts, it is typically only conducted within the first four years (Bubac et al., 2019), and importantly, inadequate monitoring may contribute to the failure of large carnivore reintroductions, such as Asiatic lions in India (Negi, 1965) and cheetah in Matusadona National Park, Zimbabwe (van der Meer et al., 2021). Large carnivore reintroductions must align with species management plans as well as local and regional policy, as programs which are unable to support self-sustaining populations through connectivity should form part of a species metapopulation plan, as demonstrated in southern African large carnivore studies (Buk et al., 2018; Davies-Mostert et al., 2015; Miller & Funston, 2014). While such strategic planning can aid the reintroduction process, as in the Dinaric lynx population establishment hunting ban (Sindičić et al., 2010), such policies are arguably most critical in managing and linking growing and stable populations. Community acceptance, tolerance and support, effective monitoring, threat intervention, disease control and veterinary support are considered important indicators, particularly during the release, establishment, and reproduction stages, while conservation strategies, action plans and policy are considered important during growth and stability stages.

#### Prioritizing stage-dependent indicators

Understandably, restoration efforts are often resource-constrained (e.g., finances, infrastructure, and expertise) and therefore, when necessary, we propose a triage approach to monitoring and reporting, such that evidence of indicators is prioritized within stages to evaluate success (Figure 3). The likelihood of project failure was thus considered for each indicator at each stage of a particular reintroduction, as derived from reviewed studies. Here we underline high priority monitoring of indicators at their respective stages: movement indicators (i.e., homing behavior as well as home range stabilization and consistency) were considered of high priority post-release until the population establishes within the release area, whereafter the importance of such monitoring decreases. Monitoring habitat, diet, sociality (i.e., intra-and inter-specific social dynamics), and resilience sociality (i.e., mortality and survival) indicators are a high priority post-release until reproduction, thereafter, monitoring importance decreases. Naturally, the breeding indicator is considered a high priority within reproduction stage, while the population growth indicator is most important in the growth stage. Sustainability indicators (i.e., dispersal and range stability, carrying capacity, population viability, and genetic diversity) are considered of highest priority within the growth and stability stages. Whereas management-released indicators (i.e., community support, effective monitoring, threat intervention, and veterinary support) are considered of high priority post-release until the reproduction stage, while larger landscape strategies, action plans and policy indicators are considered of highest priority throughout the growth and stability stages.

**FIGURE 3.**
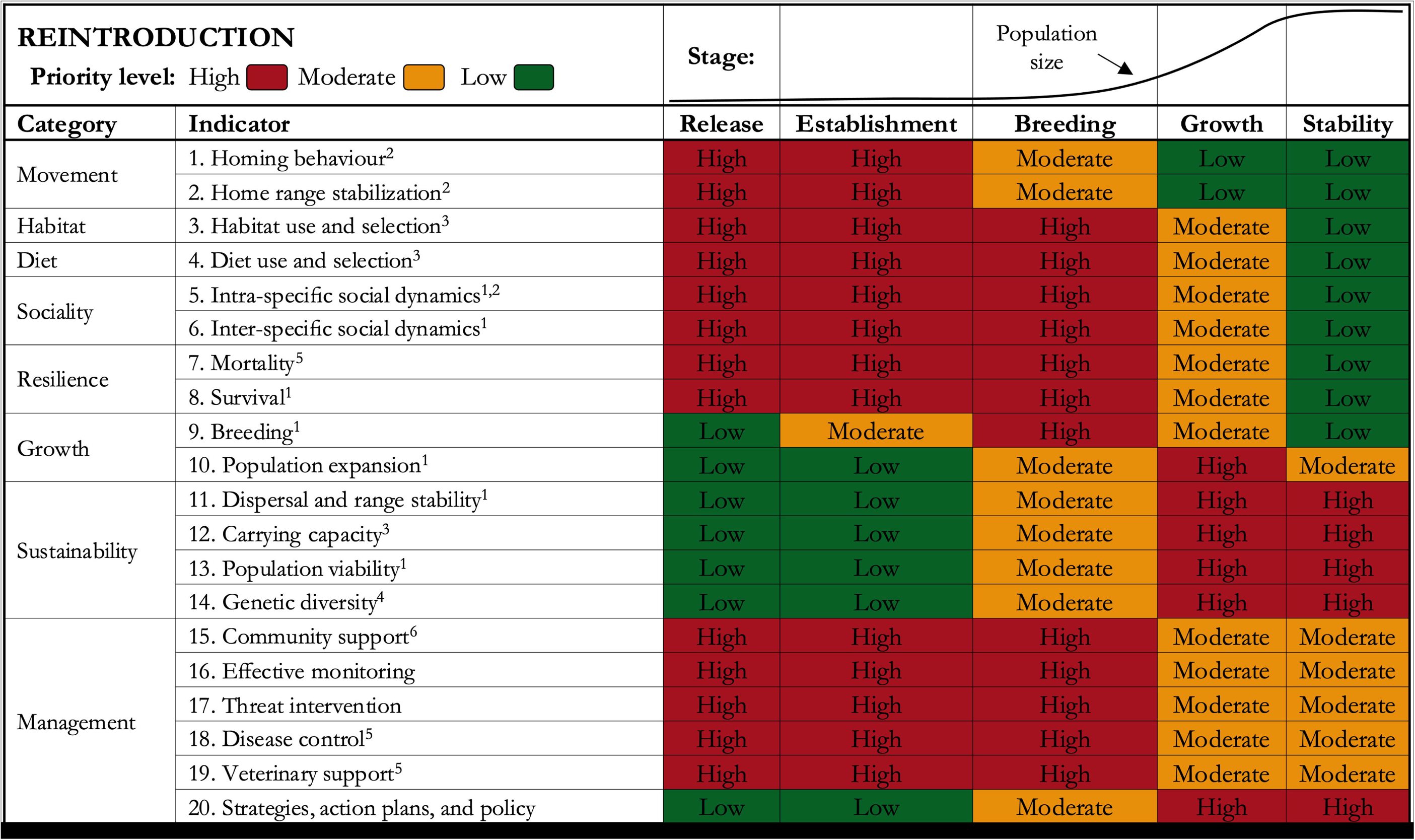
Proposed review-derived quantifiable indicators for evaluating large carnivore reintroduction success prioritized by stage within IUCN/SSC (2013) guidelines: demographic performance^1^, behavioral monitoring^2^, ecological monitoring^3^, genetic monitoring^4^, health and mortality monitoring^5^, and social, cultural and economic monitoring^6^. Using a triage approach, priorities were assigned as: i) high, ii) moderate, and iii) low likelihood of this indicator resulting in a reintroduction failure if unevaluated and the target is not met. This stage-based indicator framework should be considered a sliding scale of transitional priorities.

### Phase III: Scoring and gap analyses

Effective reintroduction evaluation requires indicators to be analysed by specific and, where applicable, quantifiable targets to determine success or failure (Miller et al., 2014). We thus proposed clearly defined, review-derived, and standardized data types, methods, and analyses required to evaluate targets for each of the 20 indicators (Table 4). We then developed an objective scoring tool based on mean target-evaluation (i.e., evaluating whether indicator-based targets were met) and indicator-coverage comparability metrics weighted by stage priority (Table S3). The reintroduction scores (%) thus emphasise both high-priority target monitoring and indicator comparability, where outcomes were defined as: ‘complete success’ (≥95% of indicators quantified and targets met), ‘success’ (75–94%), ‘partial success’ (60–74%), ‘indiscernible’ (40– 59%), ‘partial failure’ (25–39%), ‘failure’ (5–24%), and ‘complete failure’ (<5%), where success (≥75%) is considered ‘relative’ for all transitional stages, except stability, where the hand-over of a stable reintroduced population to conventional management would constitute an ‘overall’ reintroduction success. We applied this scoring system to three previously reviewed reintroductions as representative case studies to demonstrate the utility of this tool in the continuous evaluation of current and future reintroduction efforts (Figure 4). While this this scoring system could retrospectively be applied across all reviewed reintroduction cases, this would be inappropriate, given that these have collectively been used to develop our suite of indicators and that the objective was not to criticize past studies given novelty of this framework.

**FIGURE 4.**
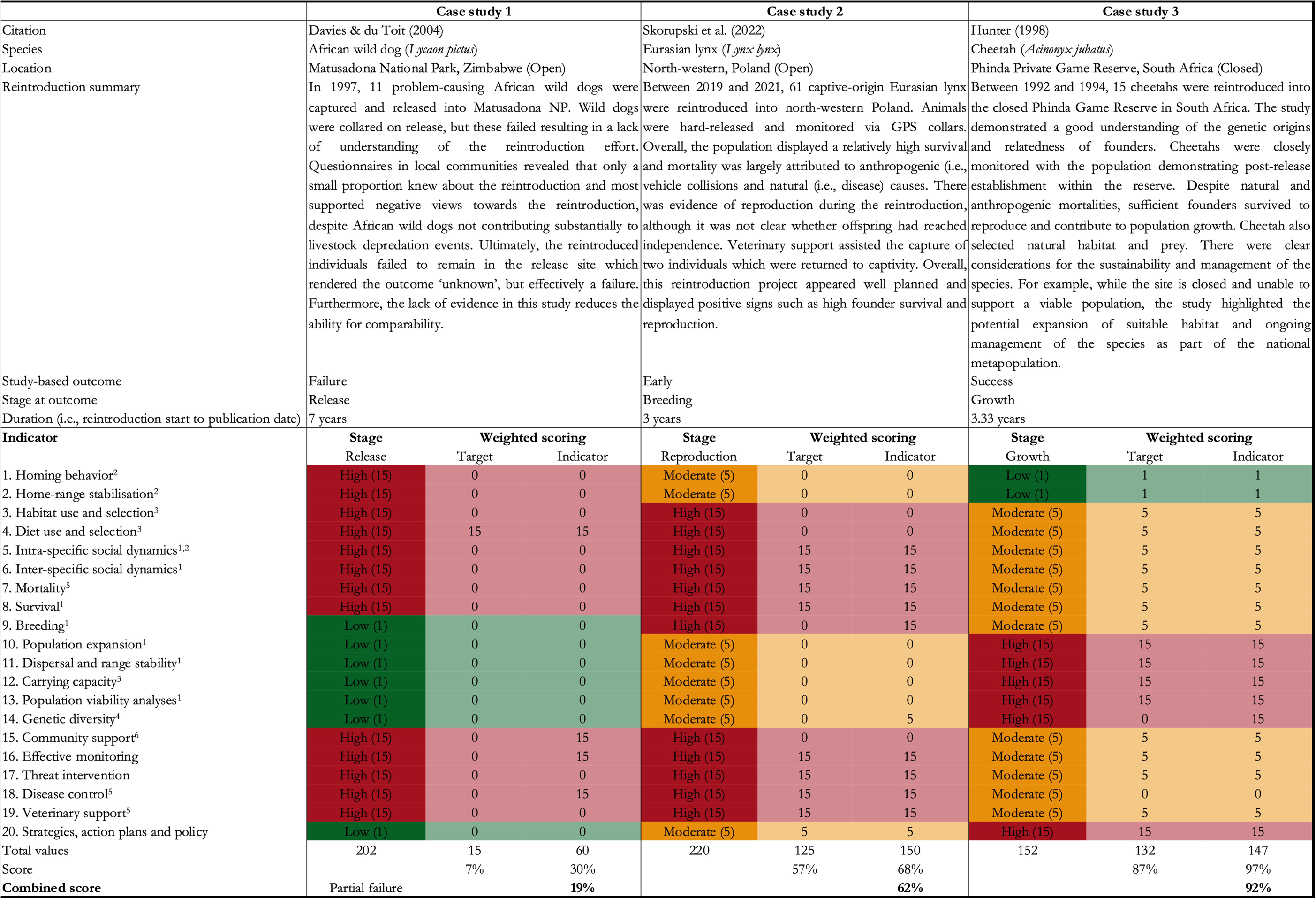
Reintroduction evaluation scoring tool applied to three reviewed reintroductions as representative case studies selected for their outcome range representation. While it is inappropriate to apply this scoring system to the reviewed reintroduction cases in this study, given that these have collectively been used to develop our suite of indicators, we did conceptually apply such scoring to three representative case studies, illustrating how such stage-depended priority scoring could be used as a tool to continuously evaluate current and future reintroduction efforts. Indicated are the study references, species, location, a brief summary of the case, Stepkovitch outcome, stage at outcome determination and study duration. Following the stage-dependent evaluation scoring tool developed in this study (Table S3), priority weighted indicators and targets were scored for each indicator and summarized as a combined overall score and category for each of the three representative case studies.

**TABLE 4.**
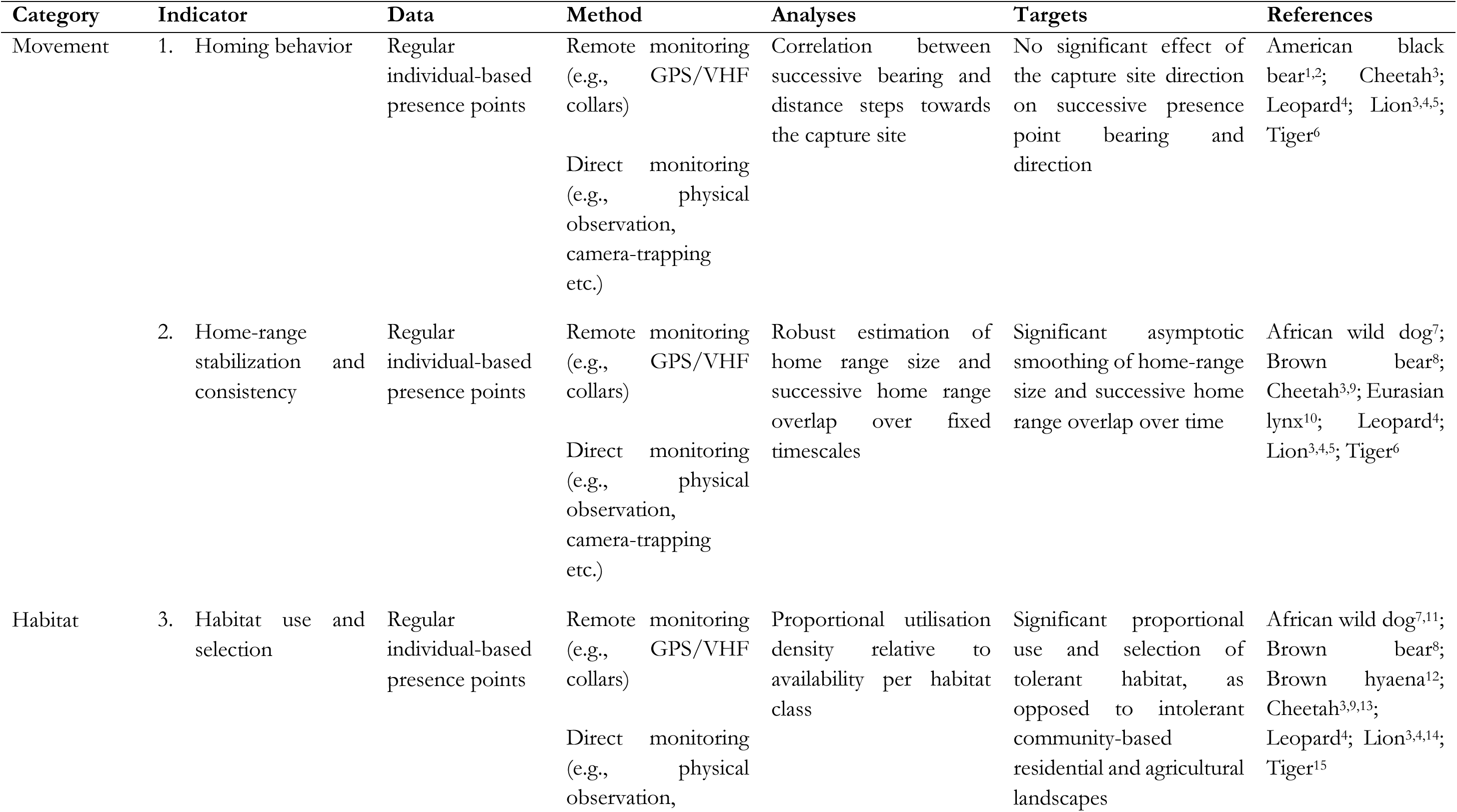
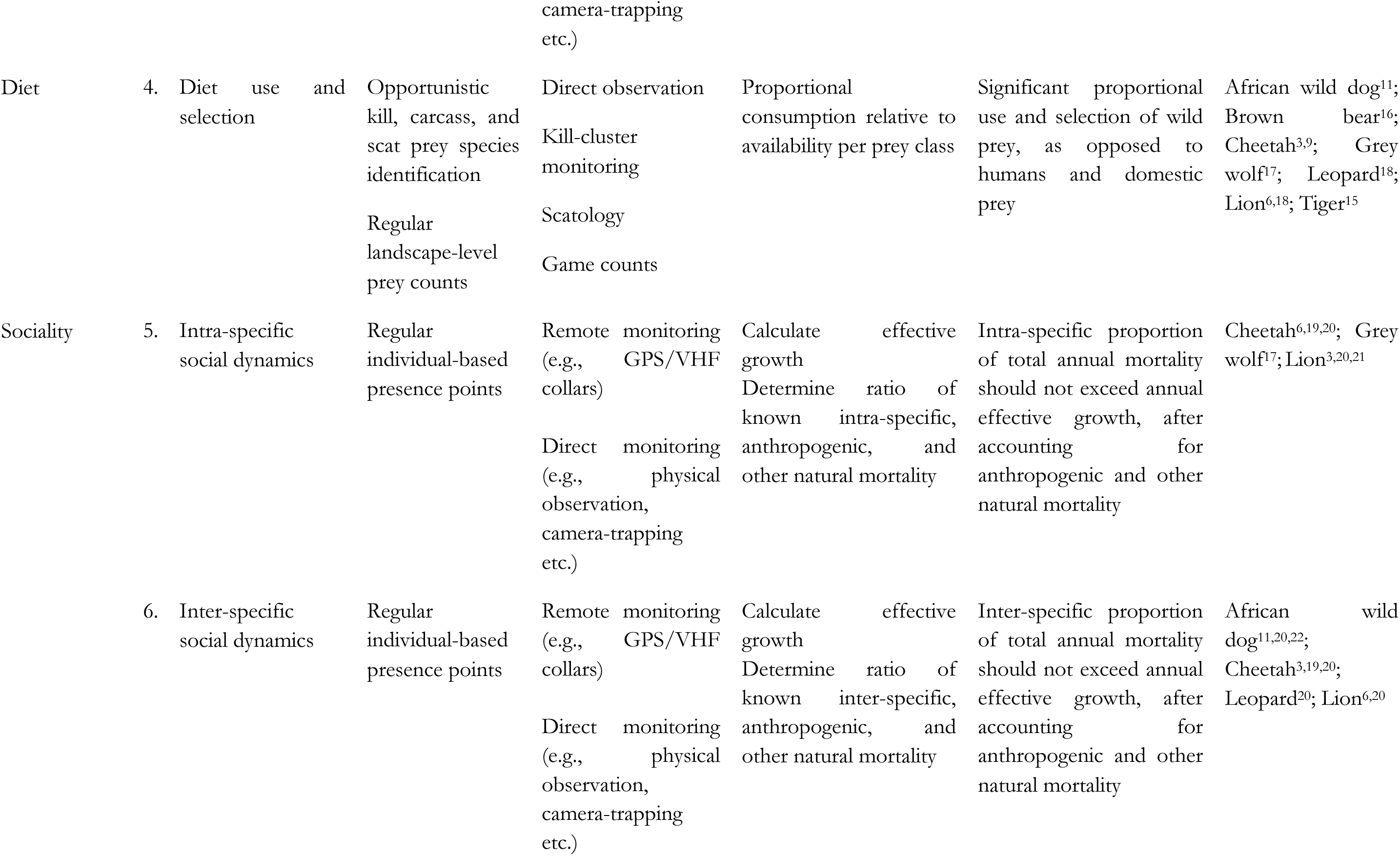
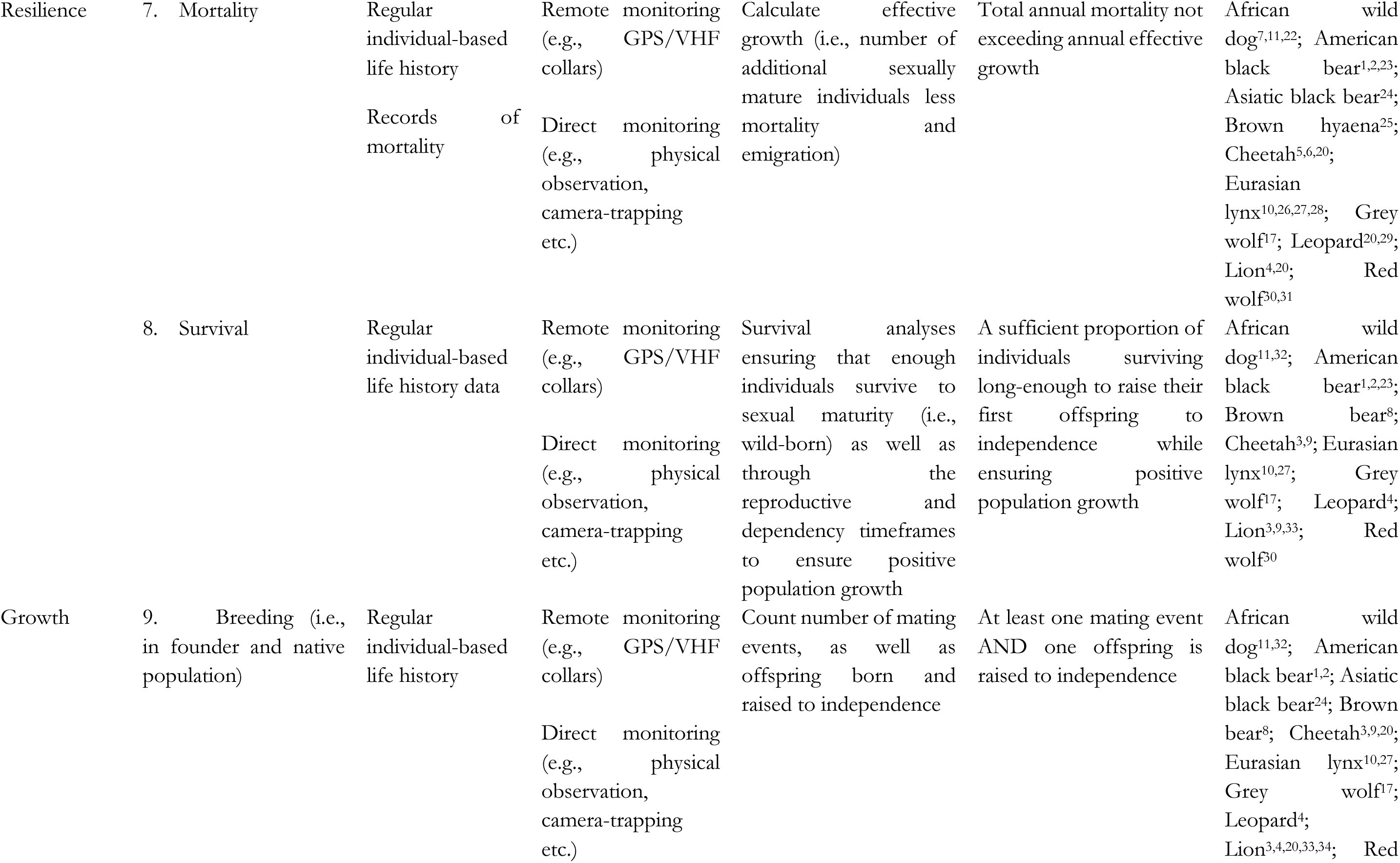
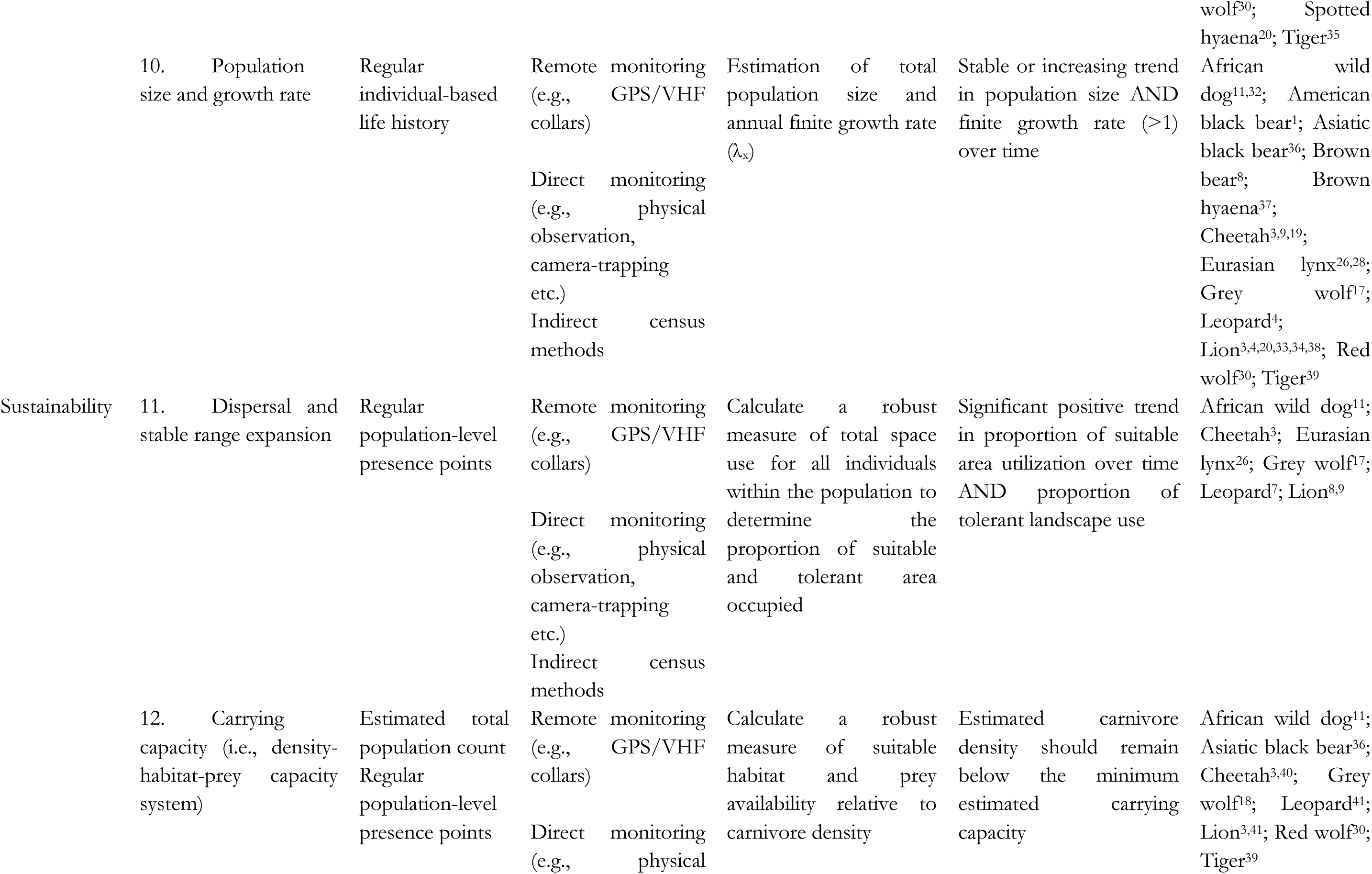
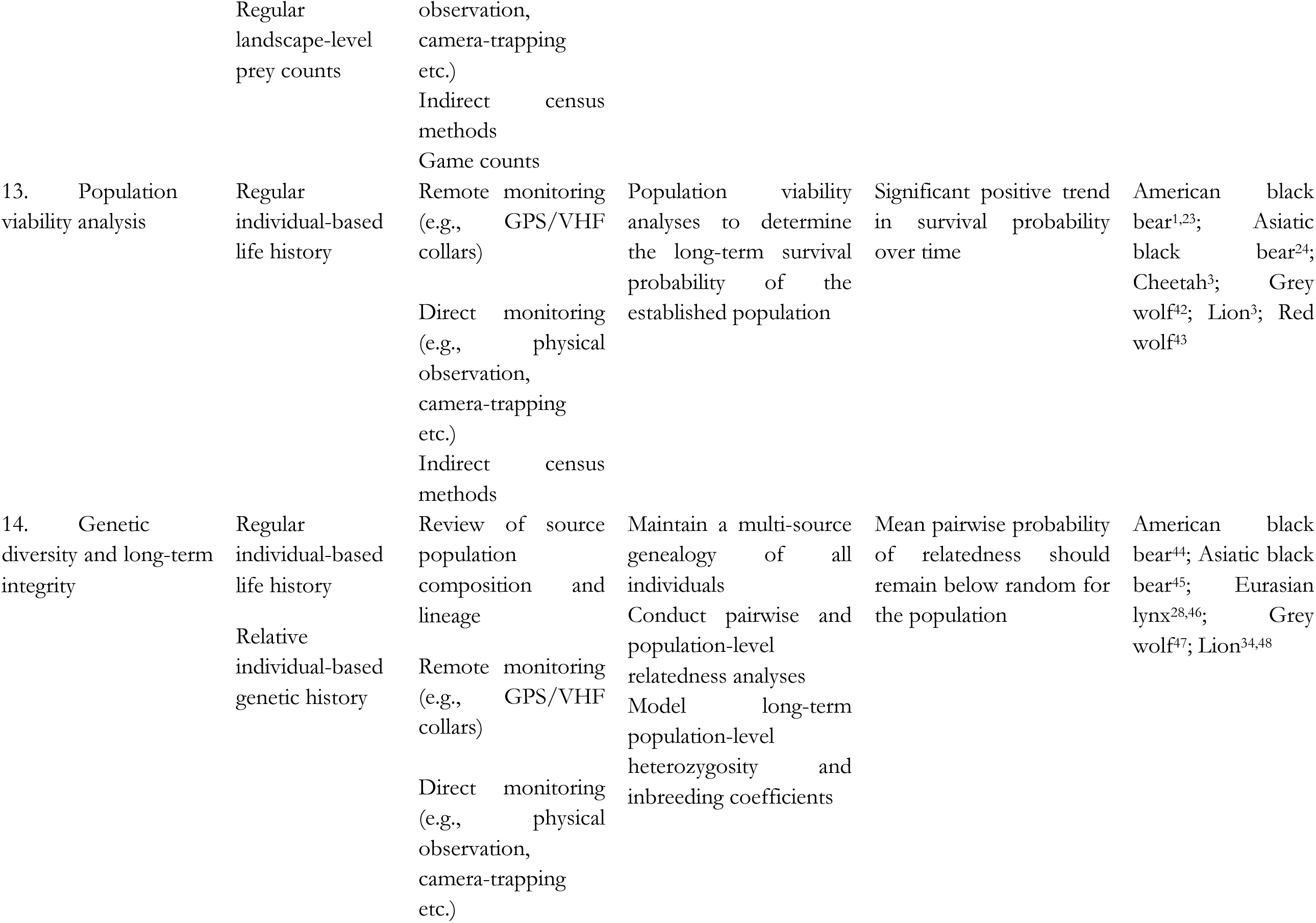
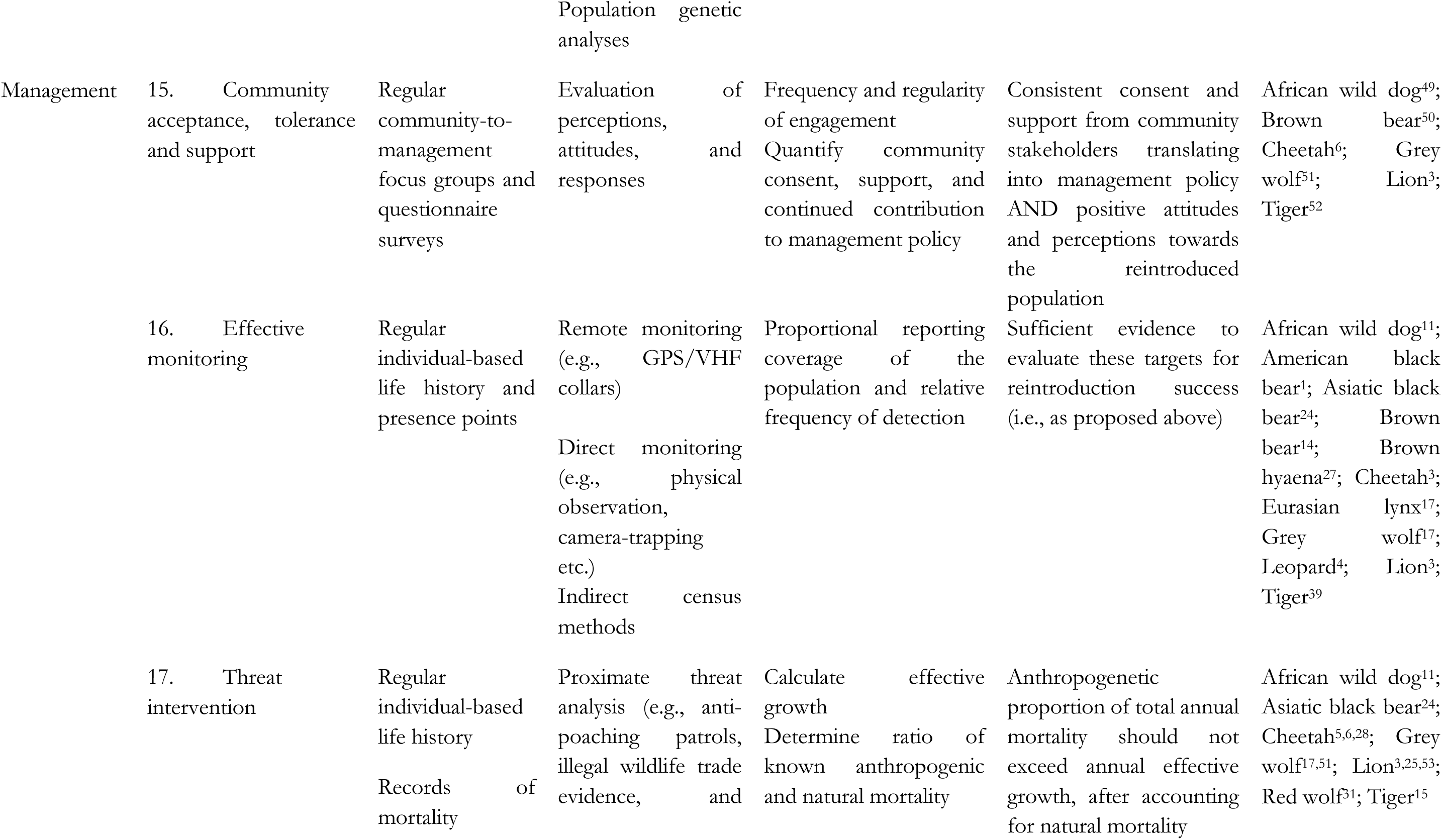
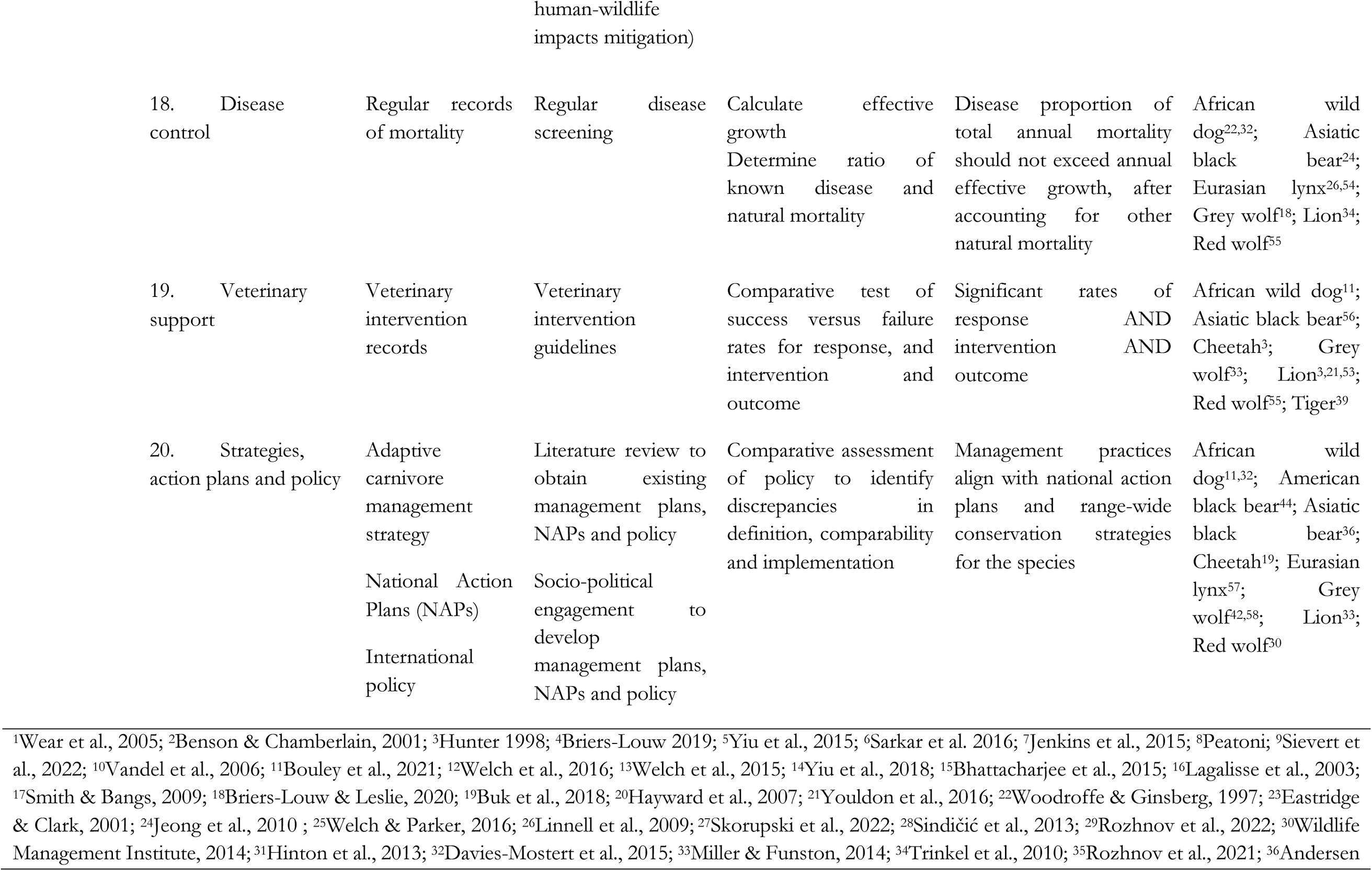
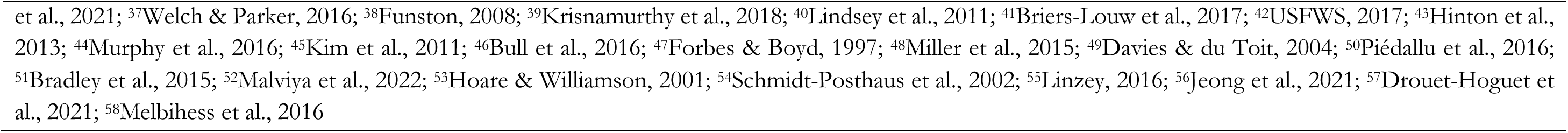
Specific and measurable targets, including data type, method, and analyses, for comparably evaluating the relative success of large carnivore reintroductions.

Instead, based on retrospective knowledge gap analysis of reviewed studies using the proposed stages (Table 2) and indicators (Table 3), we found that on average only 32% of studies quantified high priority indicators across reintroduction stages (Figure 5). The average percentage of evaluated indicators at high priority stages were as follows: release (15%), establishment (36%), breeding (36%), growth (42%), and stability (48%). Indicators frequently evaluated at high priority included: population expansion (92%), reproduction (90%), effective monitoring (55–100%), home range stabilization (i.e., 86% at establishment stage), strategies, action plans and policy (i.e., 81% at stability stage), and survival (i.e., 70% at reproduction stage). Other indicators were rarely evaluated at high priority stages, including community support (0–20%), threat intervention (0–23%), disease control (8–14%), intra-(8–15%) and inter-specific social dynamics (14–15%), and population viability (13–25%).

**FIGURE 5.**
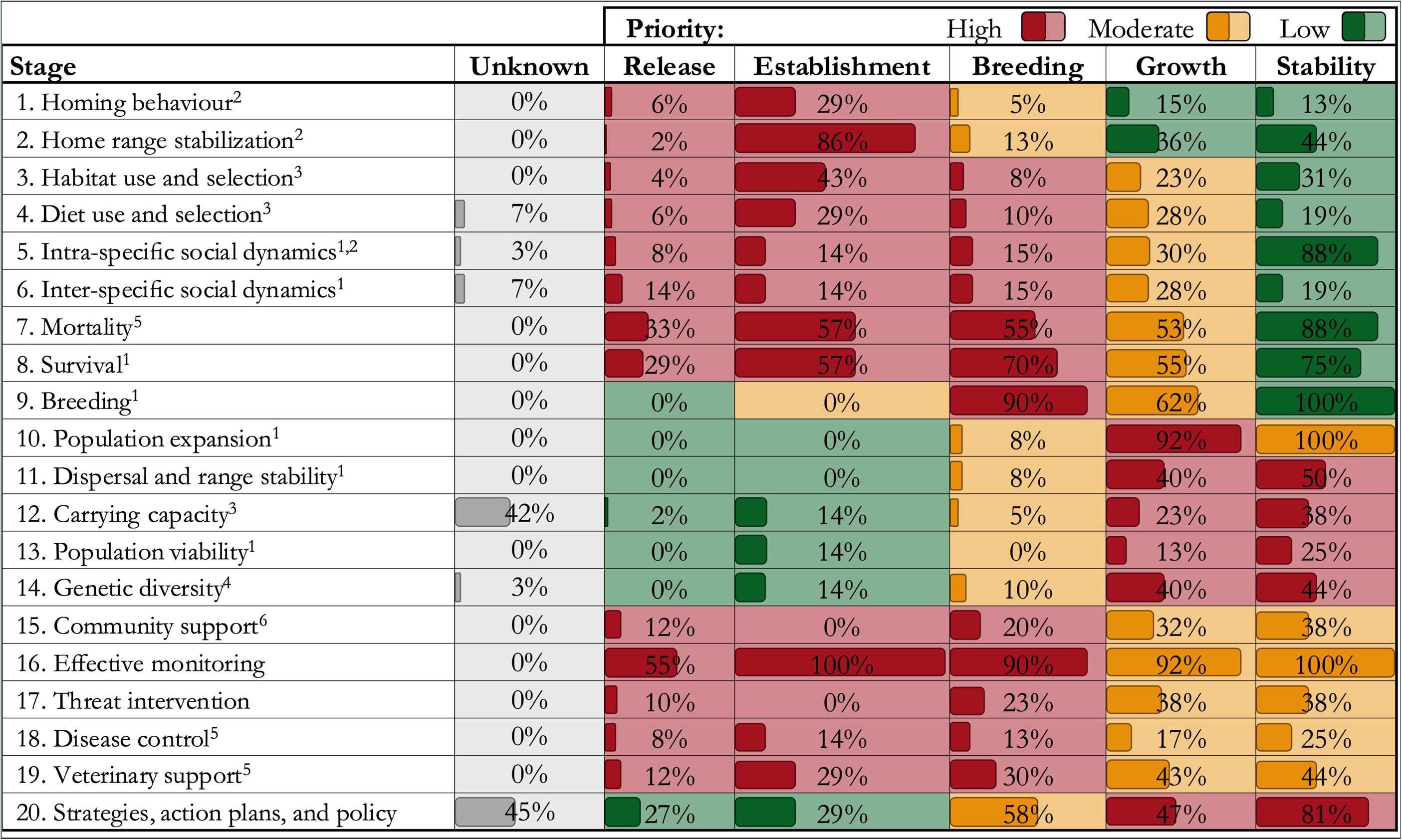
Retrospective gap analysis of reviewed large carnivore reintroduction studies (*n* = 227) relative to proposed stage-specific indicator prioritization, expressed as percentages (%). Indicators are prioritized by stage within IUCN/SSC (2013) guidelines: demographic performance^1^, behavioral monitoring^2^, ecological monitoring^3^, genetic monitoring^4^, health and mortality monitoring^5^, and social, cultural and economic monitoring

## DISCUSSION

Reintroductions provide a potential solution to the growing challenge of anthropogenic pressures which drive large carnivore population declines and extirpations worldwide (Ripple et al., 2014). However, the lack of a standardized evaluation framework hinders the efficacy, replicability, and comparability of such efforts (Morris et al., 2021). Given that sufficient information from available literature documenting over a century of reintroduction efforts now exists, we reviewed 227 large carnivore reintroductions. The aim was to contextualize past efforts and develop a comprehensive framework for evaluating reintroductions of large carnivores, extracting a full suite of clearly defined stages, indicators, and quantifiable targets. Our proposed framework addresses this current knowledge gap by improving our ability to objectively compare reintroduction efforts across programs, species and landscapes, with the support of a comprehensive management tool for the effective reintroduction of threatened species.

Based on our review, 47% of reintroduced large carnivore species were classified within ‘Threatened’ categories, of which 60% are experiencing global population declines (IUCN, 2023). Evidently, threatened status appears an important consideration in large carnivore reintroductions, although other factors such as ecotourism or restoring ecological functionality may drive reintroductions locally (Hayward et al., 2007a; Stepkovitch et al., 2022). A large percentage of reintroduction events were of Felidae (70%), which is likely due to their charismatic and iconic status (Macdonald et al., 2022). Geographically, the majority of reviewed reintroductions occurred in southern Africa (70%). This is largely explained by the transition from livestock farming to wildlife-based management practices since the 1990s particularly in South Africa, resulting in a substantial increase of small, closed reserves managed for conservation, which create adequate conditions to reintroduce large carnivores due to minimized negative human-carnivore interactions (Gusset et al., 2007; Hayward et al., 2007a; Marnewick et al., 2009). While these reintroductions were generally considered successful, relatively small systems cannot support self-sustaining populations, and thus require ongoing metapopulation management (e.g., African wild dogs, Davies-Mostert et al., 2015; cheetahs, Buk et al., 2018; and lions, Miller et al., 2013). The rest of Africa, however, mostly comprises of large, open landscapes with potential to support self-sustaining populations provided human-wildlife interactions are effectively addressed. While these landscapes typically lack funding and resources to initiate and effectively monitor such reintroductions (Lindsey et al., 2018), collaborative management partnerships between NGOs and wildlife authorities, which typically attract significant investment and enable improved management, law enforcement and infrastructure, have increased the capacity for large carnivore reintroductions (Lindsey et al., 2021). These areas are likely to become the focal point of future efforts – and indeed several reintroductions have occurred in such contexts in recent years. Arguably the most successful reintroductions within large, open landscapes include American black bears and grey wolves in North America, which have likely attained self-sustaining levels (Clark et al., 2002; Smith & Bangs, 2009). Meanwhile, human pressures have suppressed several reintroduced species, most notably certain Eurasian lynx populations across Europe due to poaching (Bull et al., 2016). Recent efforts to reintroduce jaguars in Argentina (Alberts, 2021), Persian leopards in Western Caucasus (van Gelder, 2023), and Amur leopards in the Russian Far East (Rozhnov et al., 2022) further illustrate the widespread use of this restoration tool and emphasize the need for comparability across reintroduction efforts through a unified framework.

Our review, framework, scoring tool, and retrospective gap analysis highlight relatively widespread ambiguity of reporting reintroduction success in available literature. Reintroductions were comprehensively evaluated in only a few studies (e.g., Bouley et al., 2021), with as little as 36% of cases evaluating five or more high priority indicators. Of particular concern is that 21% (*n* = 48) of reviewed cases were assigned outcomes without effective evaluation of any high priority indicators, despite sufficient time to do so, highlighting a lack of knowledge transfer and the inability to verify whether these reintroductions were actually successful. While large carnivores inherently have slow life histories, with reintroduction stages often taking several years to transition, protracted evaluations reduce the efficiency of comparability. With our priority-based framework, we hope to bridge this gap and facilitate reintroduction evaluations in real-time at defined and transitory stages throughout the reintroduction process. Furthermore, based on the retrospective analysis, only one third of studies quantified indicators at high priority stages, suggesting that two thirds of reviewed studies did not make information publicly available or contained insufficient information to effectively evaluate reintroductions at their respective stages. Despite a decrease in reintroduction failures over the past century, success rates have not improved, while the number of unevaluated ‘early’ and ‘unknown’ outcomes have increased (Stepkovitch et al., 2022), further emphasising the need for improved knowledge transfer. This reiterates the findings of Bubac et al. (2019), where most monitoring studies were conducted in the first four years post-release, which has been identified as a critical period, given the high likelihood of failure. Importantly, rather than criticize or re-evaluate reintroduction efforts, the retrospective gap analysis helps to highlight the inconsistency and lack of standardization in reintroduction efforts, while the proposed framework provides clearly defined indicators and targets to reduce vague interpretations which have significant ramifications.

While our framework provides imperative guidelines for the evaluation of future large carnivore reintroductions, we acknowledge potential shortfalls associated with oversimplification and site-or species-specific nuances. For example, in contrast to another review on large carnivore translocations (Thomas et al., 2023), where translocation success was significantly higher in unfenced compared to fenced areas, large carnivore reintroductions in our review demonstrated greater success in closed (69%) compared to open areas (31%). Additionally, the complexities associated with release area ecology and management (e.g., habitat features, prey availability, predators densities, reserve size, threat levels, and community tolerance) are difficult to account for across studies (Hayward et al., 2007a). Inter-and intra-specific species life history and ecological variability present further challenges in comparative assessments, and certain species may be more difficult to reintroduce compared to others. For example, African wild dogs and grey wolves exhibit social dependency which strongly affect social demographic structures (Gusset et al., 2006; Hurford et al., 2006), cheetahs are vulnerable to predation from larger-bodied carnivores due to their morphological differences relative to guild members (Buk et al., 2018), leopards and brown hyaenas are relatively difficult to monitor due to their secretive and elusive nature (e.g., Hayward et al., 2007b), American black bears are prone to homing (Eastridge & Clark, 2001), and red wolves suffer from hybridization with coyotes (*Canis latrans*), a small gene pool, and significantly reduced habitat (<1% remaining) and therefore require ongoing management and rewilding of captive-raised animals (Hedrick & Fredrickson, 2008). Nevertheless, we argue that the framework is sufficiently general for applicability across at least 30 large carnivores (Table 1). We note that the framework is not entirely prescriptive, but rather foundational, and may thus require modifications, particularly when applied across different taxonomic groups (e.g., smaller carnivores). Additionally, we acknowledge that not all reintroduction efforts may have been captured, while in some cases, outcomes could not be determined (i.e., unknown), although the lack of information further emphasizes the need for meticulous documentation of reintroduction efforts. Nevertheless, to advance reintroduction science, it is essential that we learn from the past successes and perhaps more so from failures, using these cases as opportunities to improve future restoration efforts by making definitive headway towards enhanced practice and policy of threatened species (Miller et al., 2014).

Holistic ecosystem recovery is only complete with sustainable top-down regulatory mechanisms which are often the most challenging to achieve when practitioners are unable to transfer their experiences and knowledge between programs, species, and landscapes (Heinen et al., 2020). Admittedly, the reintroduction of large carnivores does not necessarily translate into immediate ecosystem restoration (Alston et al., 2019). However, regulation of prey and mesocarnivore populations contribute towards reversing degraded ecosystems (Wallach et al., 2015) and the conservation of these charismatic and flagship large carnivores may benefit numerous species indirectly (Verschueren et al., 2024). Our moral obligation to restore these ecologically significant species within their former ranges should be matched by an obligation to maximize conservation value, contribute to knowledge systems, and take responsibility when such efforts are ineffective, limiting future failures which may compromise the recovery of threatened species and their ecosystems. The growing reality of large carnivore extirpations worldwide, coupled with the increased rate of reintroductions over the past century, suggest that this management tool will play an increasingly important role in future restoration efforts. For example, support for our proposed framework may immediately and directly benefit scheduled reintroductions of lions in West Africa (Aglissi et al., 2023), tigers in Asia (e.g., Chestin et al., 2017), clouded leopard (*Neofelis nebulosa*) in Taiwan (Greenspan et al., 2020), Eurasian lynx in the United Kingdom (Drouilly & O’Riain, 2021), and giant otter (*Pteronura brasiliensis*) in Argentina (Zamboni et al., 2017). While our framework applies directly to large carnivores, we urge policymakers to consider this framework for broader-scale applications. Improving the transparency and scalability of restoration efforts will help garner long-term stakeholder support, where practitioners and policy-makers should learn from failures and promote best-practices (Berger-Tal et al., 2020). Finally, incorporation of this framework into future reintroductions will effectively contribute towards reintroduction science and improve faunal rewilding efforts, aiding global ecosystem restoration.

## Ethical Statement

This research abided by the Code of Ethics of the Society for Conservation Biology. The research design did not require ethical approval from the principal investigator’s institution.

## Data Availability Statement

Supplementary files will be made available online.

## ACKNOWLEDGEMENTS

We acknowledge Zambeze Delta Conservation for supporting some of the authors.

## REFERENCES

1. Aglissi, J., Sogbohossou, E. A., & Bauer, H. (2023). Community perspectives on the prospect of lion (*Panthera leo*) reintroduction to Comoé National Park, Côte d’Ivoire (West Africa). *Wildlife Biology*, e01116.

2. Alberts, E. C. (2021). Big cat comeback: jaguars prowl Argentina’s Iberá Wetlands after 70 years. Available at: https://news.mongabay.com/2021/01/big-cat-comeback-jaguars-prowl-argentinas-ibera-wetlands-after-70-years/

3. Alston, J., Maitland, B. M., Brito, B., Esmaeili, S., Ford, A., Hays, B., Jesmer, B., Molina, F., & Goheen, J. (2019). Reciprocity in restoration ecology: when might large carnivore reintroduction restore ecosystems? Biological Conservation, 234, 82–89.

4. Arlettaz, R., Chapron, G., Kéry, M., Klaus, E., Mettaz, S., Roder, S., Vignali, S., Zimmermann, F., & Braunisch, V. (2021). Poaching threatens the establishment of a lynx population, highlighting the need for a centralized judiciary approach. Frontiers in Conservation Science, 2, 665000.

5. Athreya, V., Odden, M., Linnell, J. D. C., & Karanth, K. U. (2011). Translocation as a tool for mitigating conflict with leopards in human-dominated landscapes of India. Conservation Biology, 25, 133–141.

6. Becker, M. S., Almeida, J., Begg, C., Bertola, L., Breitenmoser, C., Breitenmoser, U., Coals, P., Funston, P., Gaylard, A., Groom, R., Henschel, P., Ikanda, D., Jorge, A., Kruger, J., Lindsey, P., Maimbo, H., Mandisodza-Chikerema, R., Maude, G., Mwape, H., et al. (2022). Guidelines for evaluating the conservation value of African lion (*Panthera leo*) translocations. Frontiers in Conservation Science, 3, 963961.

7. Berger-Tal, O., Blumstein, D. T., & Swaisgood, R. R. (2020). Conservation translocations: a review of common difficulties and promising directions. Animal Conservation, 23, 121–131.

8. Bouley, P., Paulo, A., Angela, M., du Plessis, C., & Marneweck, D. (2021). The successful reintroduction of African wild dogs (*Lycaon pictus*) to Gorongosa National Park, Mozambique. PLoS ONE, 16, e0249860.

9. Briers-Louw, W. D., & Leslie, A. J. (2020). Dietary partitioning of three large carnivores in Majete Wildlife Reserve. African Journal of Ecology, 58, 371–382.

10. Briers-Louw, W. D., Verschueren, S., & Leslie, A. J. (2019). Big cats return to Majete Wildlife Reserve, Malawi: evaluating reintroduction success. African Journal of Wildlife Research, 49, 34–50.

11. Bubac, C. M., Johnson, A., Fox, J., & Cullingham, C. (2019). Conservation translocations and post-release monitoring: identifying trends in failures, biases, and challenges from around the world. Biological Conservation, 238, 108239.

12. Buk, K., van der Merwe, V., Marnewick, K., & Funston, P. (2018). Conservation of severely fragmented populations: lessons from the transformation of uncoordinated reintroductions of cheetahs (*Acinonyx jubatus*) into a managed metapopulation with self-sustained growth. Biodiversity and Conservation, 27, 3393–3423.

13. Bull, J. K., Heurich, M., Saveljev, A. P., Schmidt, K., Fickel, J., & Förster, D. W. (2016). The effect of reintroductions on the genetic variability in Eurasian lynx populations: the cases of Bohemian-Bavarian and Vosges-Palatinian populations. Conservation Genetics, 17, 1229–1234.

14. Ceballos, G., Ehrlich, P. R., & Raven, P. H. (2020). Vertebrates on the brink as indicators of biological annihilation and the sixth mass extinction. PNAS, 117, 13596–13602.

15. Chestin, I. E., Paltsyn, M. Y., Pereladova, O. B., Iegorova, L. V., & Gibbs, J. P. (2017). Tiger re-establishment potential to former Caspaian tiger (*Panthera tigris virgata*) range in Central Asia. Biological Conservation, 205, 42–51.

16. Clark, J. D., Huber, D., & Servheen, C. (2002). Bear reintroductions: Lessons and challenges. Ursus, 13, 335–345.

17. Conservation Measures Partnership (2020). Open standards for the practice of conservation. Version 4.0. Available at: https://conservationstandards.org/wp-content/uploads/sites/3/2020/10/CMP-Open-Standards-for-the-Practice-of-Conservation-v4.0.pdf

18. Cornhill, K. L., & Kerley, G. I. H. (2020). Does competition shape cheetah prey use following African wild dog reintroductions? African Journal of Wildlife Research, 50, 75–85.

19. Cusack, J. J., Bradfer-Lawrence, T., Baynham-Herd, Z., Castelló y Tickell, S., Duporge, I., Hegre, H., Zárate, L. M., Naude, V. N., Nijhawan, J., Wilson, J., Cortes, D. G. Z., & Bunnefeld, N. (2020). Measuring the intensity of conflicts in conservation. Conservation Letters, 14, e12783.

20. Davies-Mostert, H. T, Mills, M. G. L., & Macdonald, D. W. (2009). “A critical assessment of South Africa’s managed metapopulation recovery strategy for African wild dogs,” in Reintroduction of top-order predators, eds. M. W. Hayward., and M. J. Somers (Oxford, UK: Wiley-Blackwell), 10-42.

21. Davies-Mostert, H. T., Mills, M. G. L., & Macdonald, D.W. (2015). The demography and dynamics of an expanding, managed Africa wild dog metapopulation. African Journal of Wildlife Research, 45, 258–273.

22. Drouilly, M., & O’Riain, M. J. (2021). Rewilding the world’s large carnivores without neglecting the human dimension: A response to reintroducing the Eurasian lynx to southern Scotland, England and Wales. Biodiversity and Conservation, 30, 917–923.

23. du Preez, B., Hart, T., Loveridge, A. J., & Macdonald, D. W. (2015). Impact of risk on animal behaviour and habitat transition probabilities. Animal Behaviour, 100, 22–37.

24. Eastridge, R., & Clark, J. (2001). Evaluation of 2 soft-release techniques to reintroduce black bears. Wildlife Society Bulletin, 29, 1163–1174.

25. Fox, J., & Hong, J. (2009). Effect displays in R for multinomial and proportional-odds logit models: Extensions to the effects package. Journal of Statistical Software, 32, 1–24.

26. Fox, J., & Weisberg, S. (2011). An R Companion to Applied Regression. Sage, Thousand Oaks, CA, USA.

27. Greenspan, E., Giordano, A. J., Nielsen, C. K., Sun, N. C-M., & Pei, K. J-C. (2020). Evaluating support for clouded leopard reintroduction in Taiwan: insights from surveys of indigenous and urban communities. Human Ecology, 48, 733–747.

28. Gusset, M., Slotow, R., & Somers, M. J. (2006). Divided we fail: the importance of social integration for the re-introduction of endangered wild dogs (*Lycaon pictus*). Journal of Zoology, 270, 502–511.

29. Gusset, M., Ryan, S. J., Hofmeyr, M., van Dyk, G., Davies-Mostert, H. T., Graf, J. A., Owen, C., Szykman, M., Macdonald, D. W., Monfort, S. L., Wildt, D. E., Maddock, A. H., Mills, M. G. L., Slotow, R., & Somers, M. J. (2007). Efforts going to the dogs? Evaluating attempts to re-introduce endangered wild dogs in South Africa. Journal of Applied Ecology, 45, 100–108.

30. Hale, R., Blumstein, D. T., MacNally, R., & Swearer, S. E. (2020). Harnessing knowledge of animal behavior to improve habitat restoration outcomes. Ecosphere, 11 (4), e03104.

31. Harzing, A-W., & Alakangas, S. (2016). Google Scholar, Scopus and the Web of Science: a longitudinal and cross-disciplinary comparison. Scientometrics, 106, 787–804.

32. Hayward, M. W., Adendorff, J., O’Brien, J., Sholto-Douglas, A., Bissett, C., Moolman, L. C., Bean, P., Fogarty, A., Howarth, D., Slater, R., & Kerley, G. I. H. (2007a). Practical considerations for the reintroduction of large, terrestrial, mammalian predators based on reintroductions to South Africa’s Eastern Cape Province. The Open Conservation Biology Journal, 1, 1–11.

33. Hayward, M. W., Adendorff, J., O’Brien, J., Sholto-Douglas, A., Bissett, C., Moolman, L. C., Bean, P., Fogarty, A., Howarth, D., Slater, R., & Kerley, G. I. H. (2007b). The reintroduction of large carnivores to the Eastern Cape, South Africa: an assessment. Oryx, 41, 205–214.

34. Hedrick, P. W., & Fredrickson, R. J. (2008). Captive breeding and the reintroduction of Mexican and red wolves. Molecular Ecology, 17, 344–350.

35. Heinen, J. H., Rahbek, C., & Borregaard, M. K. (2020). Conservation of species interactions to achieve self-sustaining ecosystems. Ecography, 43, 1603–1611.

36. Heurich, M., Schultze-Naumburg, J., Piacenza, N., Magg, N., Červeny, J., Engleder, T., Heerdtfelder, M., Sladova, M., & Kramer-Schadt, S. (2018). Illegal hunting as a major driver of the source-sink dynamics of a reintroduced lynx population in Central Europe. Biological Conservation, 224, 355–365.

37. Hinton, J. W., Brzeski, K. E., Rabon Jr, D. R., & Chamberlain, M. J. (2015). Effects of anthropogenic mortality on critically endangered red wolf *Canis rufus* breeding pairs: implications for red wolf recovery. Oryx, 51, 174–181.

38. Hunter, L. T. (1998). The behavioural ecology of reintroduced lions and cheetahs in the Phinda Resource Reserve, KwaZulu-Natal, South Africa. PhD thesis, University of Pretoria, Pretoria, South Africa.

39. Hunter, L. T. B., Pretorius, K., Carlisle, L. C., Rickelton, M., Walker, C., Slotow, R., & Skinner, J. D. (2007). Restoring lions Panthera leo to northern KwaZulu-Natal, South Africa: short-term bio-logical and technical success but equivocal long-term conservation. Oryx, 41, 196–204.

40. Hurford, A., Hebblewhite, M., & Lewis, M. A. (2006). A spatially explicit model for an Allee effect: why wolves recolonize so slowly in Greater Yellowstone. Theoretical Population Biology, 70, 244–154.

41. IUCN/SSC (2013). Guidelines for Reintroductions and Other Conservation Translocations. Version 1.0. IUCN Species Survival Commission, Gland, Switzerland.

42. IUCN (2023). IUCN Red List of Threatened Species. IUCN Global Species Programme Red List Unit, Cambridge, United Kingdom. http://iucnredlist.org

43. Jenkins, E., Silva-Opps, M., Opps, S. B., & Perrin, M. (2015). Home range and habitat selection of a reintroduced African wild dog (*Lycaon pictus*) pack in a small South African game reserve. South African Journal of Wildlife Research, 45, 233–246.

44. Jeong, D-H., Jang, K., Tang, J-J., Choi, J.Y., Lim, S-H., Yeon, S-C., Shim, K. M., Kim, S. E., & Kang, S. S. (2021). Treatment of two Asiatic black bears (*Ursus thibetanus*) with severe injuries and their subsequent release into the wild: a case report. BMC Veterinary Research, 17, 125.

45. Jepson, P. R. (2022). To capitalise on the decade of ecosystem restoration, we need institutional redesign to empower advances in restoration ecology and rewilding. People and Nature, 4, 1404–1413.

46. Jimenez, M. D., Bangs, E. E., Boyd, D. K., Smith, D. W., Becker, S. A., Ausband, D. E., Woodruff, S. P., Bradley, E. H., Holyan, J., & Laudon, K. (2017). Wolf dispersal in the Rocky Mountains, Western United States: 1993–2008. Journal of Wildlife Management, 81, 581–592.

47. Kolipaka, S. S., Tamis, W. L. M., van ‘t Zelfde, M., Persoon, G. A., & de Iongh, H. H. (2017). Wild versus domestic prey in the diet of reintroduced tigers (*Panthera tigris*) in the livestock-dominated multiple-use forests of Panna Tiger Reserve, India. PLoS ONE, 12, e0174844.

48. Lenth, R. V. (2016). Least-squares means: The R package lsmeans. Journal of Statistical Software, 69, 1–33.

49. Lindsey, P., Tambling, C. J., Brummer, R., Davies-Mostert, H., Hayward, M., Marnewick. K., & Parker, D. (2011). Minimum prey and area requirements of the vulnerable cheetah *Acinonyx jubatus*: implications for reintroduction and management of the species in South Africa. Oryx, 45, 587–599.

50. Lindsey, P., Baghai, M., Bigurube, G., Cunliffe, S., Dickman, A., Fitzgerald, K., Flyman, M., Gandiwa, P., Kumchedwa, B., Madope, A., Morjan, M., Parker, A., Steiner, K., Tumenta, P., Uiseb, K., & Robson, A. (2021). Attracting investment for Africa’s protected areas by creating enabling environments for collaborative management partnerships. Biological Conservation, 255, 108979.

51. Lindsey, P. A., Miller, J. R. B., Petracca, L. S., Coad, L., Dickman, A. J., Fitzgerald, K. H., Flyman, M. V., Funston, P. J., Henschel, P., Kasiki, S., Knights, K., Loveridge, A. J., MacDonald, D. W., Mandisodza-Chikerema, R. L., Nazerali, S., Plumptre, A. J., Stevens, R., Van Zyl, H. W., & Hunter, L. T. B. (2018). More than $1 billion needed annually to secure Africa’s protected areas with lions. Proceedings of the National Academy of Sciences, 115, E10788–E10796.

52. Linzey, D. W. (2016). Mammals of Great Smoky Mountains National Park: 2016 Revision. Southeastern Naturalist, 15, 1–93.

53. Macdonald, D. W., Johnson, P. J., Burnham, D., Dickman, A., Hinks, A., Sillero-Zubiri, C., & Macdonald, E. A. (2022). Understanding nuanced preferences for carnivore conservation: to know them is not always to love them. Global Ecology and Conservation, 37, e02150.

54. Mahli, Y., Franklin, J., Seddon, N., Solan, M., Turner, M. G., Field, C. B., & Knowlton, N. (2020). Climate change and ecosystems: threats, opportunities and solutions. Philosophical Transactions of the Royal Society B, 375, 20190104.

55. Malviya, M., Kalyanasundaram, S., & Krishnamurthy, R. (2022). Paradox of success-mediated conflicts: analysing attitudes of local communities towards successfully reintroduced tigers in India. Frontiers in Conservation Science, 2, 783467.

56. Marnewick, K., Hayward, M. W., Cilliers, D., & Somers, M. J. (2009). Survival of cheetahs relocated from ranchland to fenced protected areas in South Africa. In: Hayward MW, Somers MJ (eds) Reintroduction of top-order predators, 282–306. Wiley-Blackwell, Oxford, UK.

57. Miller, K. A., Bell, T. P., & Germano, J. M. (2014). Understanding publication bias in reintroduction biology by assessing translocations of New Zealand’s herpetofauna. Conservation Biology, 28, 1045–1056.

58. Miller, S. M., Bissett, C., Burger, A., Courtenay, B., Dickerson, T., Druce, D.J., Ferreira, S., Funston, P. J., Hofmeyr, D., Kilian, P. J., Matthews, W., Naylor, S., Parker, D. M., Slotow, R., Toft, M., & Zimmermann, D. (2013). Management of reintroduced lions in small, fenced reserves in South Africa: an assessment and guidelines. South African Journal of Wildlife Research, 43, 138–154.

59. Miller, S. M., & Funston, P. J. (2014). Rapid growth rates of lion (*Panthera leo*) populations in small, fenced reserves in South Africa: a management dilemma. South African Journal of Wildlife Research, 44, 43–55.

60. Morris, S. D., Brook, B. W., Moseby, K. E., & Johnson, C. N. (2021). Factors affecting success of conservation translocations of terrestrial vertebrates: a global systematic review. Global Ecology and Conservation, 28, e01630.

61. Muhly, T. B., & Musiani, M. (2009). Livestock depredation by wolves and the ranching economy in the Northwestern U.S. Ecological Economics, 68, 2439–2450.

62. Murphy, S. M., Cox, J. J., Augustine, B. C., Hast, J. T., Guthrie, J. M., Wright, J., McDermott, J., Maehr, S. C., & Plaxico, J. H. (2016). Characterizing recolonization by a reintroduced bear population using genetic spatial capture-recapture. Journal of Wildlife Management, 80, 1390–1407.

63. Negi, S. S. (1965). Transplanting of Indian lion in Uttar Pradesh state. Cheetal, 12, 98–101.

64. Pettorelli, N., Barlow, J., Stephens, P. A., Durant, S. M., Connor, B., Bühne, H. S., Sandom, C. J., Wentworth, J., & du Toit, J. T. (2018). Making rewilding fit for policy. Journal of Applied Ecology, 55, 1114–1125.

65. Phiri, C. (1996). Cheetah translocation project in Lower Zambezi National Park, Zambia. National Parks and Wildlife Service, Zambia.

66. Preatoni, D., Mustoni, A. Martinoli, A., Carlini, E., Chiarenzi, B., Chiozzini, S., Van Dongen, S., Wauters, L. A., & Tosi, G. (2005). Conservation of brown bear in the Alps: space use and settlement behavior of reintroduced bears. Acta Oecologica, 28, 189–197.

67. Purchase, G. (2007). Mozambique: preliminary assessment of the status and distribution of cheetah. CAT News, 3, 37–39.

68. R Core Team. (2023). R: A language and environment for statistical computing.

69. Redpath, S. M., Bhatia, S., & Young, J. (2015). Tilting at wildlife: reconsidering human-wildlife conflict. Oryx, 49, 222–225.

70. Ripple, W. J., Estes, J. A., Beschta, R. L., Wilmers, C. C., Ritchie, E. G., Hebblewhite, M., Berger, J., Elmhagen, B., Letnic, M., Nelson, M. P., Schmitz, O. J., Smith, D. W., Wallach, A. D., & Wirsing, A. J. (2014). Status and ecological effects of the world’s largest carnivores. Science, 343, 1241484.

71. Robert, A., Colas, B., Guigon, I., Kerbirou, C., Mihoub, J. B., Saint-Jalme, M., & Sarrazin, F. (2015). Defining reintroduction success using IUCN criteria for threatened species: a demographic assessment. Animal Conservation, 18, 397–406.

72. Rozhnov, V. V., Naidenko, S. V., Hernandez-Blanco, J. A., Chistopolova, M. D., Sorokin, P. A., Yachmennikova, A. A., Blidchenko, E. Y., Kalinin, A. Y., & Kastrikin, V. A. (2021). Restoration of the Amur tiger (*Panthera tigris altaica*) population in the Northwest of its distribution area. Biology Bulletin, 48, 1401–1423.

73. Rozhnov, V. V., Yachmennikova, A. A., Dronova, N. A., Naidenko, S. V., Hernandez-Blanco, J. A., Chistopolova, M. D., Pkhitikov, A. B., Tembotoba, F. A., Trepet, S. A., & Chestin, I. E. (2022). Experience of the leopard recovering through reintroduction in the Russian Caucasus. CAT News, 15, 67–71.

74. Sankar, K., Qureshi, Q., Nigam, P., Malik, P. K., Sinha, P. R., Mehrotra, R. N., Gopal, R., Bhattacharjee, S., Mondal, K., & Gupta, S. (2010). Monitoring of reintroduced tigers in Sariska Tiger Reserve, Western India: preliminary findings on home range, prey selection and food habits. Tropical Conservation Science, 3, 301–318.

75. Sarkar, M. S., Ramesh, K., Johnson, J. A., Sen, S., Nigam, P., Gupta, S. K., Murthy, R. S., & Saha, G. K. (2016). Movement and home range characteristics of reintroduced tiger (*Panthera tigris*) population in Panna Tiger Reserve, central India. European Journal of Wildlife Research, 62, 537–547.

76. Scheepers, J. L., & Venzke, K. A. E. (1995). Attempts to reintroduce African wild dogs *Lycaon pictus* into Etosha National Park, Namibia. South African Journal of Wildlife Research, 25, 138–140.

77. Schmidt-Posthaus, H., Breitenmoser-Würsten, C., Posthaus, H., Bacciarini, L., & Breitenmoser, U. (2002). Causes of mortality in reintroduced Eurasian lynx in Switzerland. Journal of Wildlife Diseases, 38, 84– 92.

78. Seddon, P. (1999). Persistence without intervention: assessing success in wildlife reintroductions. Trends in Ecology and Evolution, 14, 503.

79. Seddon, P. J., Armstrong, D. P., & Maloney, R.F. (2007). Developing the science of reintroduction biology. Conservation Biology, 21, 303–312.

80. Skorupski, J., Tracz, M., Tracz, M., & Śmietana, P. (2022). Assessment of Eurasian lynx reintroduction success and mortality risk in north-west Poland. Scientific Reports, 12, 12366.

81. Sindičić, M., Sinanović, N., Majić, S. A., Huber, D., Kunovac, S., & Kos, I. (2010). Legal status and management of the Dinaric lynx population. Veterinaria, 58, 229–238.

82. Smith, D. W., & Bangs, E. E. (2009). Reintroduction of wolves to Yellowstone National Park: history, values and ecosystem restoration. In: Hayward MW, Somers MJ (eds) Reintroduction of top-order predators, 92–125. Wiley-Blackwell, Oxford, UK.

83. Stepkovitch, B., Kingsford, R. T., & Moseby, K. E. (2022). A comprehensive review of mammalian carnivore translocations. Mammal Review, 52, 554–572.

84. Tilman, D., Clark, M., Williams, D. R., Kimmel, K., Polasky, S., & Packer, C. (2017). Future threats to biodiversity and pathways to their prevention. Nature, 546, 73–81.

85. Thomas, S., van der Merwe, V., Carvalho, W. D., Adania, C. H., Černe, R., Gomerčić, T., Krofel, M., Thompson, J., McBride Jr, R. T., Hernandez-Blanco, J., Yachmennikova, A., Macdonald, D. W., & Farhadinia, M. S. (2023). Evaluating the performance of conservation translocations in large carnivores across the world. Biological Conservation, 279, 109909.

86. Tosi, G., Chirichella, R., Zibordi, F., Mustoni, A., Giovannini, R., Groff, C., Zanin, M., & Apollonio, M. (2015). Brown bear reintroduction in the Southern Alps: To what extent are expectations being met? Journal of Nature Conservation, 26, 9–19.

87. Trinkel, M., Ferguson, N., Reid, A., Reid, C., Somers, M., Turelli, L., Graf, J., Szykman, M., Cooper, D., Haverman, P., Kastberger, G., Packer, C., & Slotow, R. (2008). Translocating lions into an inbred lion population in the Hluhluwe-iMfolozi Park, South Africa. Animal Conservation, 11, 138–143.

88. van Der Meer, E., Sousa, L. L., & Loveridge, A. J. (2021). Reassessment of an introduced cheetah Acinonyx jubatus population in Matusadona National Park, Zimbabwe. Oryx, 55, 294–301.

89. van Gelder, A. (2023). One small step for 3 Amur leopards, one giant leap for the rarest big cat species. Available at: https://conservewildcats.org/2023/06/16/amur-leopard-relocation/

90. Venables, W. N., & Ripley, B. D. (2013). *Modern Applied Statistics with S-PLUS*. Springer, New York, NY, USA.

91. Verschueren, S., Bauer, H., Cristescu, B., Leirs, H., Torres-Uribe, C., & Marker, L. (2024). From popularity to preservation: large carnivore potential for ecosystem conservation. Mammal Review, Early View. https://onlinelibrary.wiley.com/doi/full/10.1111/mam.12365

92. von Holdt, B. M., Stahler, D. R., Smith, D. W., Earl, D. A., Pollinger, J. P., & Wayne, R. K. (2008). The genealogy and genetic viability of reintroduced Yellowstone grey wolves. Molecular Ecology, 17, 252– 274.

93. Wallach, A. D., Izhaki, I., Toms, J. D., Ripple, W. J., & Shanas, U. (2015). What is an apex predator? Oikos, 124, 1453–1461.

94. Wear, B. J., Eastridge, R., & Clark, J. D. (2005). Factors affecting settling, survival, and viability of black bears reintroduced to Felsenthal National Wildlife Refuge, Arkansas. Wildlife Society Bulletin, 33, 1363– 1374.

95. Weise, F. J., Lemeris Jr. J., Stratford, K. J., van Vuuren, R. J., Munro, S. J., Crawford, S. J., Marker, L. L., & Stein, A. B. (2015). A home away from home: insights from successful leopard (*Panthera pardus*) translocations. Biodiversity and Conservation, 24, 1755–1774.

96. Welch, R. J., & Parker, D. M. (2016). Brown hyaena population explosion: rapid population growth in a small, fenced system. Wildlife Research, 43, 178–187.

97. Woodroffe, R., & Ginsberg, J. (1997). The role of captive breeding and reintroduction in wild dog conservation. In: Woodroffe R, Ginsberg J, Macdonald DW (eds) The African Wild Dog: Status Survey and Conservation Action Plan, 100-111. IUCN/SSC, Switzerland.

98. Youldon, D., Monks, N., & Abell, J. (2016). Re-introduction of the African lion from a captive origin: Zambia & Zimbabwe. In: Soorae PS (ed) Global Re-introduction Perspectives, 2016: Case-Studies from Around the Globe, 153-156. IUCN/SSC Re-introduction Specialist Group, Abu Dhabi, UAE.

99. Zamboni, T., Di Martino, S., & Jiménez-Pérez, I. (2017). A review of a multispecies reintroduction to restore a large ecosystem: The Iberá Rewilding Program (Argentina). Perspectives in Ecology and Conservation, 15, 248–256.

