## Supplementary material for "Guidelines for evaluating the success of large carnivore reintroductions": Table S1

**S U P P O R T I N G	F I L E S**

**Table S1.** Definitions of outcomes, based on Stepkovitch et al. (2022), assigned to large reviewed carnivore reintroductions studies.

| Outcome | Definition |
| --- | --- |
| Failure | Translocation failed, as dictated by text. Includes mortality of some/all founders or project aims not achieved.  ^©^ Considered failure originally but population detected after  monitoring decades later.  ^+^ Originally considered successful but population locally  extinct in subsequent years or decades.  ^®^ Failure due to individuals/pack being removed from reserve  (usually due to high prey losses) |
| Partial success | Partial success, as dictated by text. I.e., Breeding observed but self-sustaining population not achieved yet and/or further supplementations required.  * Achieved experiment aims but did not necessarily lead to  recovery/establishment of self-sustaining population.  ^A^ Further augmentations required |
| Full success | Successful reintroduction or augmentation, as dictated by text. Generally breeding observed, high survivorship of founders and population still extant in subsequent surveys.  ^A^ Further augmentations required  ^®^ Initially successful but individuals/pack subsequently  removed from reserve (usually due to high prey losses)  ^ST^ Successful in short term, but ongoing monitoring required  ^M^ Management required – removal and/or culling of individuals  to manage populations reaching carrying capacity inside  reintroduction area  ^>^ Did not establish inside release area but population  established outside release area as a result of translocation  ^#^ Too successful – subjected to population control as a result of  increase in numbers  ^?^ Assumed success – no post-release monitoring but species  now common |
| Early | Translocated individuals alive, but no assessment or information on the success of the translocation. Includes projects undertaken within three years of the publication where no other post-monitoring information was available.  ^ breeding observed  ^o^ Project ongoing at time of publication  ** High mortality observed but population still extant |
| Unknown | Not specified in text or online |

**Table S2.** List of reviewed large carnivore reintroductions studies (*n* = 227). Table S2 can be found as a separate supplementary spreadsheet as it was not possible to fit within the margins of this document.

**Table S3.** Template for evaluating large carnivore reintroductions based on the scoring system. Table S3 can be found as a separate supplementary spreadsheet as it was not possible to fit within the margins of this document.
